# Scalable and universal prediction of cellular phenotypes enables in silico experiments

**DOI:** 10.1101/2024.08.12.607533

**Authors:** Yuge Ji, Alejandro Tejada-Lapuerta, Niklas A. Schmacke, Zihe Zheng, Xinyue Zhang, Simrah Khan, Ina Rothenaigner, Juliane Tschuck, Kamyar Hadian, Veit Hornung, Fabian J. Theis

## Abstract

Biological systems can be interrogated by perturbing individual components and observing the consequences across molecular, cellular, and phenotypic levels. The vast combinatorial space of possible perturbations and responses makes exhaustive experimentation infeasible. Recent advances in machine learning have shown that training on diverse datasets enables transfer learning across tasks, capturing patterns that generalize and improving performance on previously unseen problems. Inspired by this principle, we present Prophet, a transformer-based model pretrained on a vast, heterogeneous collection of perturbation experiments. This pretraining allows Prophet to predict the outcomes of untested genetic or chemical perturbations in novel cellular contexts, spanning phenotypes such as gene expression, viability, and morphology. By leveraging shared structure across apparently disconnected assays, Prophet provides a scalable framework for large-scale virtual screening and prioritization of informative experiments. Prophet consistently outperforms baseline models, including those trained on single phenotypes, showing that transfer learning between phenotypes not only is possible but improves predictive accuracy. Its capabilities extends to in vivo developmental systems, where it recapitulates known lineage biology and proposes new candidates. In a large-scale in silico screen for melanoma, Prophet identified and experimentally validated compounds with selective activity that mirrored clinically approved therapies, demonstrating its ability to transform perturbation biology into a predictive and scalable engine for therapeutic discovery.

## Introduction

High-throughput screening technologies have revolutionized cellular biology, enabling the simultaneous testing of tens of thousands of compounds or genetic perturbations. For example, small molecule and CRISPR screens have been conducted to systematically interrogate the biological mechanisms of cancer resistance ^1,2^, identify new drug targets ^3,4^, capture synergistic interactions^5^, and deconvolve drug mechanisms of action ^6–9^. Yet the immense combinatorial space of perturbations, cell states, and phenotypes makes exhaustive experimentation infeasible. Accurate in silico modeling could guide the selection of informative experiments, reduce the number of screens required for discovery ^10^, and identify cell-context specific compounds while simultaneously making findings more robust across technical variation ^11^. Existing statistical and AI-based analyses have advanced individual screens ^8,12^ but generally fail to integrate information across experiments ^13,14^.

Recent breakthroughs in deep learning suggest a way forward. Two principles, scalable architectures and pretraining on massive, heterogeneous datasets, have enabled transformative progress across domains. Models trained on broad distributions, such as transformers in natural language^15,16^, vision^17^, and proteins^18^, capture shared patterns that generalize across tasks and contexts. This transfer learning paradigm allows knowledge gained from one dataset to improve performance on tasks that were not explicitly part of training. Analogously, biological experiments often appear disconnected, yet share underlying structures across perturbations, cell types, and phenotypes. By leveraging large, heterogeneous experimental datasets, a model can capture these shared structures, enabling transfer learning that improves predictions in novel experimental contexts.

Here, we present Prophet (**Pr**edictor **o**f **phe**no**t**ypes), a computational model that leverages relationships across cellular phenotypes to perform in silico screening. Inspired by the success of transformers in solving complex problems across diverse domains, Prophet represents an experiment as the combination of three fundamental elements — the cellular state, the perturbations applied, and the phenotype measured — and learns predictive mappings across this space. Unlike most existing approaches, which are constrained to single datasets or endpoints, Prophet is pretrained on 4.7 million experiments spanning diverse phenotypes and assay types. We predict phenotypic responses (e.g. viability, gene expression profiles, T-cell proliferation, etc.) and provide uncertainty estimates, enabling risk-aware prioritization of experimental candidates. Notably, we demonstrate that learning from high-throughput viability assays improves predictions of more complex phenotypes such as cell morphology, highlighting the utility of leveraging cost-effective data to enhance performance on resource-intensive endpoints. Prophet not only outperforms baselines, including models trained on single phenotypes, but also demonstrates real-world utility. It improves the efficiency of virtual screening, identifies informative experiments with far fewer trials, and prospectively guides discovery, as shown by the successful experimental validation of compounds with selective activity against melanoma.

Prophet unifies heterogeneous perturbation datasets into a single predictive framework, enabling transfer of insights across assays, cell types, and phenotypes. This approach provides a principled strategy for navigating the combinatorial space of biological experiments, translating machine learning advances into experimental design.

## Results

### A unified framework for experimental cell biology

To enable transfer learning across perturbational datasets, we first developed a unified representation of experiments in cell biology. We conceptualize any experiment as the combination of three fundamental components: the cell state the experiment is performed in, the applied perturbation, and the measured phenotype (Fig. 1A, B). The cellular state represents the baseline cellular context in which the experiment is performed (e.g. cell line, cell type). The perturbation is the specific intervention or perturbation applied to the cells during the experiment. Commonly, these are small molecules or genetic perturbations (e.g. CRISPR/Cas9 knockouts, CRISPRi/a). The phenotype denotes the readout used to measure the effect of the perturbation, encompassing molecular phenotypes (e.g., gene expression profiles), macroscopic phenotypes (e.g., cell viability), and morphological phenotypes (e.g., Cell Painting’s numeric features). Within this framework, the outcome of an experiment is defined as the measurable effect of the perturbation on the given state (Supplementary Note 1). This abstraction allows diverse experimental modalities to be described in a consistent way, forming the foundation for a generalizable predictive model.

**Fig. 1.**
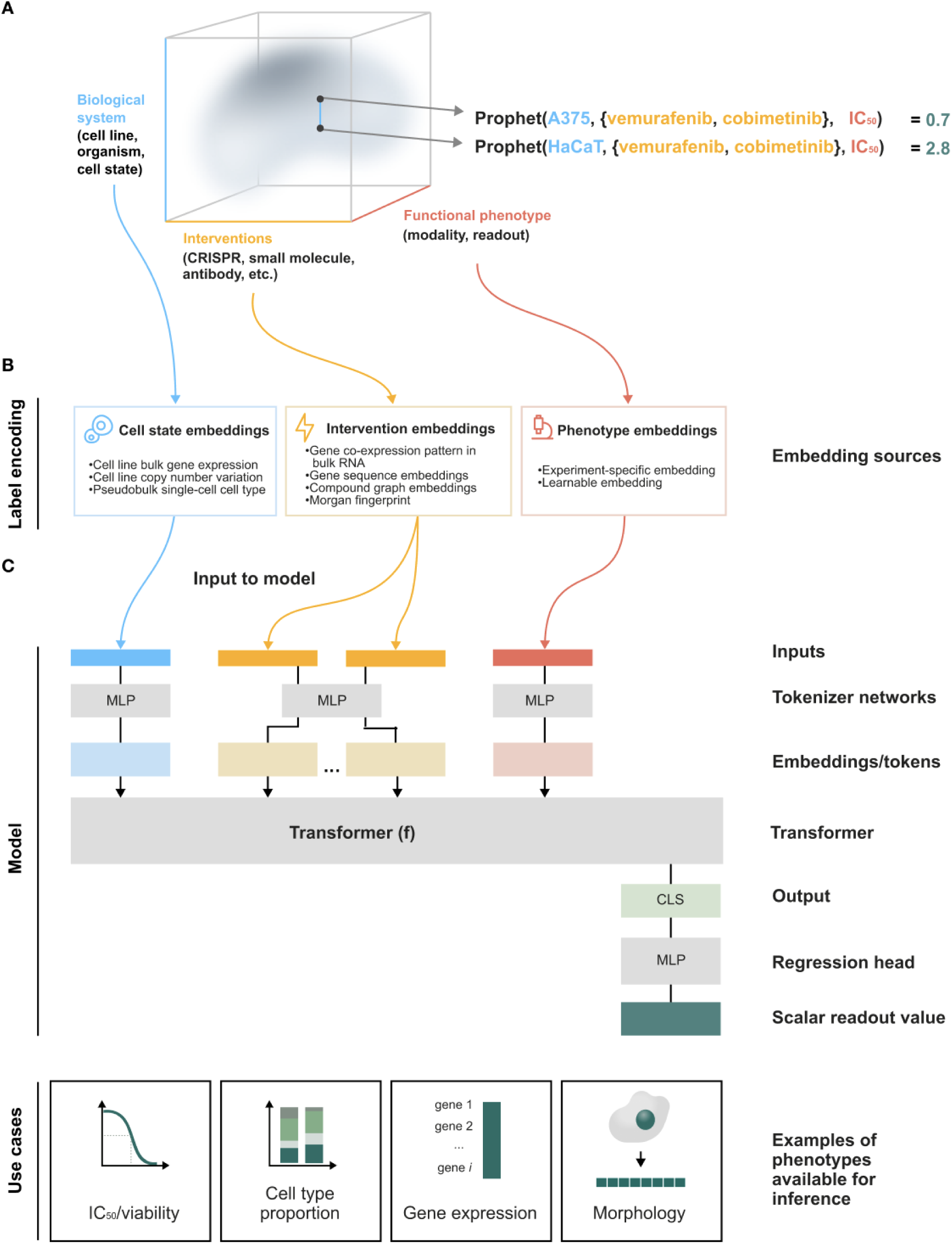
Prophet is a universal cellular phenotype predictor. (**A**) Prophet models the experimental space of cell biology along three dimensions: cellular state, perturbation, and phenotype. Each location in this 3D space corresponds to an experiment defined by a specific combination of these three factors, for example the measurement of cell viability (phenotype) in A375 cells (cellular state) treated with a chemotherapy drug (perturbation). Because this space is continuous, Prophet can model quantitative relationships between experiments and transfer knowledge across them. We illustrate one example where two experiments differ only in the cellular state they were performed in: A375 and HaCaT cells. This flexibility enables Prophet to be trained across datasets. (**B**) Cellular states, perturbations, and phenotypes are represented as vectors. These can be derived from prior knowledge (e.g., transcriptomic profiles or molecular fingerprints) or modeled as learnable embeddings, enabling flexible integration of diverse data types. (**C**) Prophet uses a transformer-based architecture with self-attention to jointly represent states, perturbations, and phenotypes. Prior knowledge is tokenized into a common embedding space, which the transformer processes to generate predictions. Prophet outputs scalar or vector-valued phenotypes, allowing prediction of diverse experimental outcomes such as viability, gene expression, or cell morphology.

To instantiate this framework, we represent each component of the experimental triplet as a vector (Fig. 1C). For cellular states, we use transcriptomic profiles from reference atlases such as the Cancer Cell Line Encyclopedia. Perturbations are encoded depending on their modality: small molecules are described by molecular fingerprints, and genetic perturbations by embeddings of their target gene’s features. Phenotypes are represented either as learnable embeddings or, when prior knowledge exists, using dedicated encoders. Importantly, this design is modular: alternative representations can be substituted as new datasets or encoding methods become available.

To train Prophet within this framework, we assembled a large compendium of perturbation-response datasets comprising ten publicly available resources: SCORE^3,19^, LINCS^20^, JUMP^8^, GDSC^3,19,21^, GDSC^22^, CTRP^23^, PRISM^1^, Horlbeck et al.^24^, Saunders et al.^25^ and Shifrut et al.^26^ (fig. S1A-B, Table S1). Each individual measurement, such as the IC50 of a drug in a given cell line, or a morphological profile (via CellPainting features) after a perturbation, is treated as a single experiment. This formulation allows Prophet to integrate highly heterogeneous datasets within a common structure.

Prophet is implemented as a supervised transformer-based model (Fig. 1C). The architecture consists of 8 transformer encoder units ^16,27^ with 8 attention heads per layer and a feed-forward network with a hidden dimensionality of 1,024 to generate a 512-dimensional embedding of each experiment (fig. S2A). Prophet leverages existing knowledge of cellular states and perturbations by projecting prior knowledge-based representations into a common token space using neural networks as tokenizers. The phenotype representations are modeled as learnable embeddings, directly projected in the token space. Through self-attention, Prophet effectively captures relationships between perturbations, cellular states, and phenotypic outcomes, facilitating nuanced learning and prediction. The output of the transformer is fed into a feed-forward network that predicts the value of the phenotype of interest as a scalar. When phenotypes are vector-valued (e.g., gene expression or Cell Painting features), Prophet decomposes them into individual scalar prediction tasks (e.g., gene-wise log-fold changes or morphological features such as cell eccentricity). Prophet is trained end-to-end using supervised learning to minimize mean square error (MSE) between the predicted and measured outcomes(fig. S2C).

This unified framework allows Prophet to integrate heterogeneous experiments within a single predictive model, providing the foundation for transfer learning across assays, perturbations, and cellular contexts.

### Transfer learning improves in silico experimental prediction

We first evaluated Prophet’s ability to predict the outcomes of unseen experiments, focusing on two biologically relevant tasks: predicting the effect of perturbations on previously untested cell states (unseen cell state), and the effect of previously untested perturbations (unseen perturbation) (fig. S1C). Predictions were benchmarked against three baseline architectures: random forest, multilayer perceptron, and linear regression, as well as the mean estimator, which summarizes the global effect of perturbations across prior experiments (fig. S1C, Methods) and is used as a difficult baseline in drug response prediction ^14^. Hit rate, coefficient of determination (R2), and Spearman correlation were used to assess prediction quality, capturing extreme responses, overall performance, and ranking accuracy, respectively (Methods).

To establish a baseline to evaluate the usefulness of transfer learning, we first trained specialized Prophet models (Prophet-individual) on each dataset individually, with the same architecture but a smaller number of parameters to prevent overfitting (Fig. 2A, Methods, Supplementary Methods, Table S2). Across all datasets, Prophet-individual consistently outperformed baselines, improving hit rates by an average of 63% over the best simpler architecture and capturing up to 33% of all hits (Fig. 2E, fig. S3A-D). Hit rates for unseen perturbations were generally higher than for unseen cell states (Fig. 2E, fig. S5C), reflecting the relative difficulty of extrapolating to new cellular contexts.

**Fig. 2.**
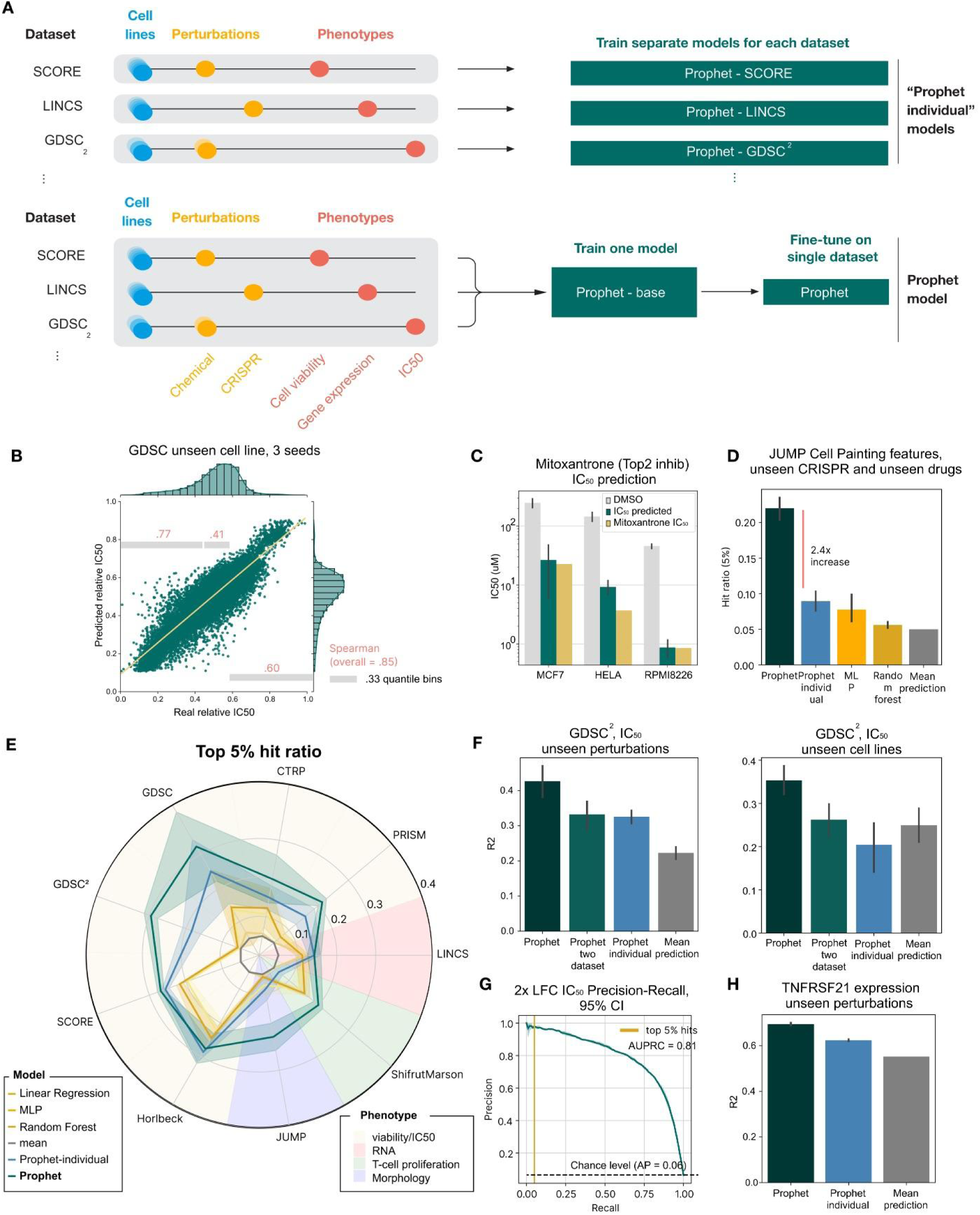
Prophet’s performance and scaling across diverse datasets. (**A**) Prophet-individual models (top) are Prophet models trained only on a single dataset. On the other hand, the full Prophet model (bottom) is pretrained using all datasets and then fine tuned on the dataset of interest. (**B**) Spearman correlation of three different bins of the data, for one split in GDSC unseen cell line prediction across three seeds. Values are binned according to the real IC50 values, with cutoffs at the ⅓ quantiles. (**C**) Prophet accurately predicts the IC_50_ value of the unseen compound mitoxantrone, a clinically approved compound absent from training data. (**D**) Performance comparison on the JUMP Cell Painting dataset for predicting unseen CRISPR and drug perturbations. Prophet significantly outperforms all baselines, achieving a 2.4× higher 5% hit ratio than Prophet individual, highlighting the benefit of transfer learning across perturbation types. (**E**) Improvement in the hit ratio score for the extrema 5% of compounds over the mean baseline of the models evaluated: linear regression, multilayer perceptron (MLP), Random Forest, single-assay Prophet (Prophet-individual) and fine-tuned Prophet (Prophet). Datasets have a background color assigned based on phenotype category, showing the diversity of prediction. (**F**) In GDSC^2^, fine-tuned Prophet outperforms Prophet-individual, the mean estimator and also a Prophet-two-dataset model trained jointly in GDSC^2^ and PRISM. Fine-tuning uses only 20,000 steps with a batch size of 256, far fewer optimization steps than other models, highlighting the effectiveness of the pretraining stage. (**G**) Precision-recall curve evaluating the ability to predict 2x fold change hits in GDSC, with shaded region indicating 95% confidence interval. The area-under the precision-recall curve (AUPRC) is 0.81, substantially exceeding the chance level (average precision = 0.06, dashed line). The vertical gold line indicates the recall corresponding to the top 5% of predicted hits, highlighting performance in the most experimentally relevant region. (**H**) Fluorescent RNA value of the gene TNFRSF21 is not part of the pre-training corpus. However, fine-tuning Prophet on it still yields better results than the mean estimator and than training a Prophet-individual model only on this dataset.

We next tested Prophet’s ability to leverage transfer learning by jointly training on two datasets (Prophet-two-dataset). We combined training on GDSC2^22^, a small molecule combinatorial perturbation viability screen reporting IC_50_ values, with PRISM^1^, a much larger single-perturbation screen with a CellTiterGlo viability phenotype. PRISM contains 8 of the 61 compounds and 76 of the 110 cell lines in GDSC2. Joint training improved prediction of unseen perturbations and cell states in GDSC2 by 3% and 28%, respectively (Fig. 2F), indicating that incorporating additional datasets can enhance performance.

Building on this principle, we pretrained our largest Prophet model on 4.7 million experimental conditions, covering 26,000 perturbations, 1,550 cell states, and multiple phenotypes (IC_50_, viability and RNA expression) across seven small molecule and genetic perturbation screens (Methods, Fig. S1A-B, Table S1). Once pretrained, the model was fine-tuned on a specific dataset of interest (Fig. 2A), which consistently matched or outperformed all baseline models, including Prophet-individual models trained on single datasets, across every benchmark (Fig. 2E, Methods, Table S3). Notably, these gains extend across small-molecule and genetic perturbations, including unseen monotherapies and combinatorial perturbations, demonstrating Prophet’s ability to leverage shared structure across diverse experiments (GDSC, CTRP, PRISM, SCORE, Shifrut, JUMP, LINCS, Horlbeck et al.).

Prophet accurately predicts the readout of novel experiments in silico. For example, Mitoxantrone, a clinically approved compound indicated for multiple cancers ^28,29^ absent from training datasets was predicted de novo, and the predicted IC50 values across cancer types agreed with measurements from GDSC and literature reports ^30,31^ (Fig. 2C). Similarly, Prophet accurately predicts complex phenotypes not included in pretraining, such as cell morphology (CellPainting features) (JUMP) and transcriptomics (LINCS) (Fig. 2D, H), highlighting its ability to transfer knowledge across diverse experimental endpoints. This generalizability serves as evidence of Prophet’s ability to enable transfer learning, leveraging information across multiple screens. As an example, pretraining followed by fine-tuning on GDSC2 improved R2 by 70% for unseen perturbations and 30% for unseen cell states compared to Prophet-individual, and outperformed a model trained on PRISM and GDSC2 by 28–29% (Fig. 2F). All improvements were statistically significant (p ≤ 0.05, fig. S4), confirming that transfer learning across datasets provides substantial predictive gains.

Prophet’s transfer learning capability is not limited to phenotypes seen during the pre-training stage. We did not pre-train Prophet on any morphological measurement, but Prophet fine-tuned on JUMP Cell Painting features outperformed both the Prophet-individual model trained only on JUMP Cell Painting features and the baseline models (Fig. 2E). We observed the same behavior with unseen RNA values: for example, the mRNA abundance value of the highly differentially expressed gene TNFRSF21 from LINCS was left out from the pre-training corpus; nonetheless, the fine-tuned version of Prophet is 25% better than the baseline and 10% better than a single-dataset Prophet model trained uniquely on TNFRSF21 values (Fig. 2H). Likewise, the same pattern can be observed even with phenotypes that not only are not included in the pretraining corpus but also are measured in completely different cell states, such as T-cell proliferation. In this case, Prophet-individual is outperformed by baselines such as RF but the fine-tuned Prophet outperforms RF by 33% (Fig. 2E, fig. S4).

Beyond global responses, Prophet enables in silico prioritization of extreme responses (“hits”) (Fig. 2E, fig. S5E) for efficient experimental planning. Using a top/bottom 5% definition of hits (Methods), Prophet increased the number of predicted logFC hits in unseen GDSC cell lines by 12.7× over random selection, achieving an area-under the precision-recall curve (AUPRC) of 0.81 (Fig. 2G, Supplementary Methods, Fig. S5B,E). With a more stringent 4× fold-change threshold, Prophet reduced the number of experiments required by 60× while maintaining high precision and no false positives among the top 1% of hits. Spearman rank and R² metrics were consistent with these findings (fig. S5D), demonstrating reliable ranking of perturbation effects across conditions.

Finally, we assessed how Prophet’s predictive uncertainty and performance scale with the number of conditions observed during training (Methods). We fine-tuned models on increasingly more training data, using datasets with enough cell states (CTRP, GDSC, GDSC2, LINCS, SCORE). This revealed that confidence in predictions increased as more perturbations and cell states were incorporated (fig. S5A). Performance similarly improved with additional data (fig. S6, S7). To fully assess Prophet’s generalization capabilities, we evaluated its performance in particularly challenging scenarios designed to stress-test the model, including unseen tissue types, novel drug scaffolds, and clusters of similar drugs and cell states deliberately left out of training. Across 64 such stratified tasks, Prophet outperformed Prophet-individual and baseline models in 69% of cases, with differences in the remaining tasks being minimal (≤ 0.015 hit ratio, fig. S3). While Prophet demonstrates robust transfer learning, we do not expect it to perform optimally when the test set is extremely far out of distribution relative to the training data.

Despite its strong performance, Prophet encounters limitations in certain scenarios. For example, predicting the effects of observed perturbations in untested cell lines within SCORE, or forecasting the impact of untested small molecules in known cell lines within LINCS, can be challenging. Further analysis revealed that the correlation between experimental replicates is a key determinant of predictability (Table S4, Methods). In some cases, replicates of the same experimental condition are less similar to each other than to replicates of other random conditions, reflecting high experimental noise and batch effects. Under such circumstances, the mean estimator represents an upper bound on predictive performance.

In summary, these results demonstrate that pretraining Prophet on large, diverse experimental datasets enables effective transfer learning, improving predictions for unseen perturbations, cell states, and phenotypes, and supporting efficient in silico prioritization of experimental candidates.

### Prophet learns a universal landscape of biological experiments

Drug screens are limited in scope, either by the number of compounds tested or the variety of cell states. Hence, they provide a limited view of the profile of a compound, encompassing just the effects in the tested cell states. Including more cell states would provide a more robust and informative view of the pharmacological profile of a drug^1^. Now, with Prophet, we can examine the perturbation effect *in silico* across hundreds of cell states at once. We represent each small molecule—or combinations of small molecules—as a vector in which each dimension is the Prophet-predicted viability value in each cancer cell state (Fig. 3A). This vector represents an “activity-based fingerprint” for each compound. In this high-dimensional representation, compounds with similar sensitivity profiles across cancer cell lines cluster together when viewed using a 2D representation (Fig. 3B). We can draw comparisons between these shared response patterns for unbiased annotations of mechanisms of action (Fig. 3C), identification of new therapeutic candidates (Fig. 3D, E), or isolation of tissue-specific mechanisms (Fig. 3F, fig. S9A).

**Fig. 3.**
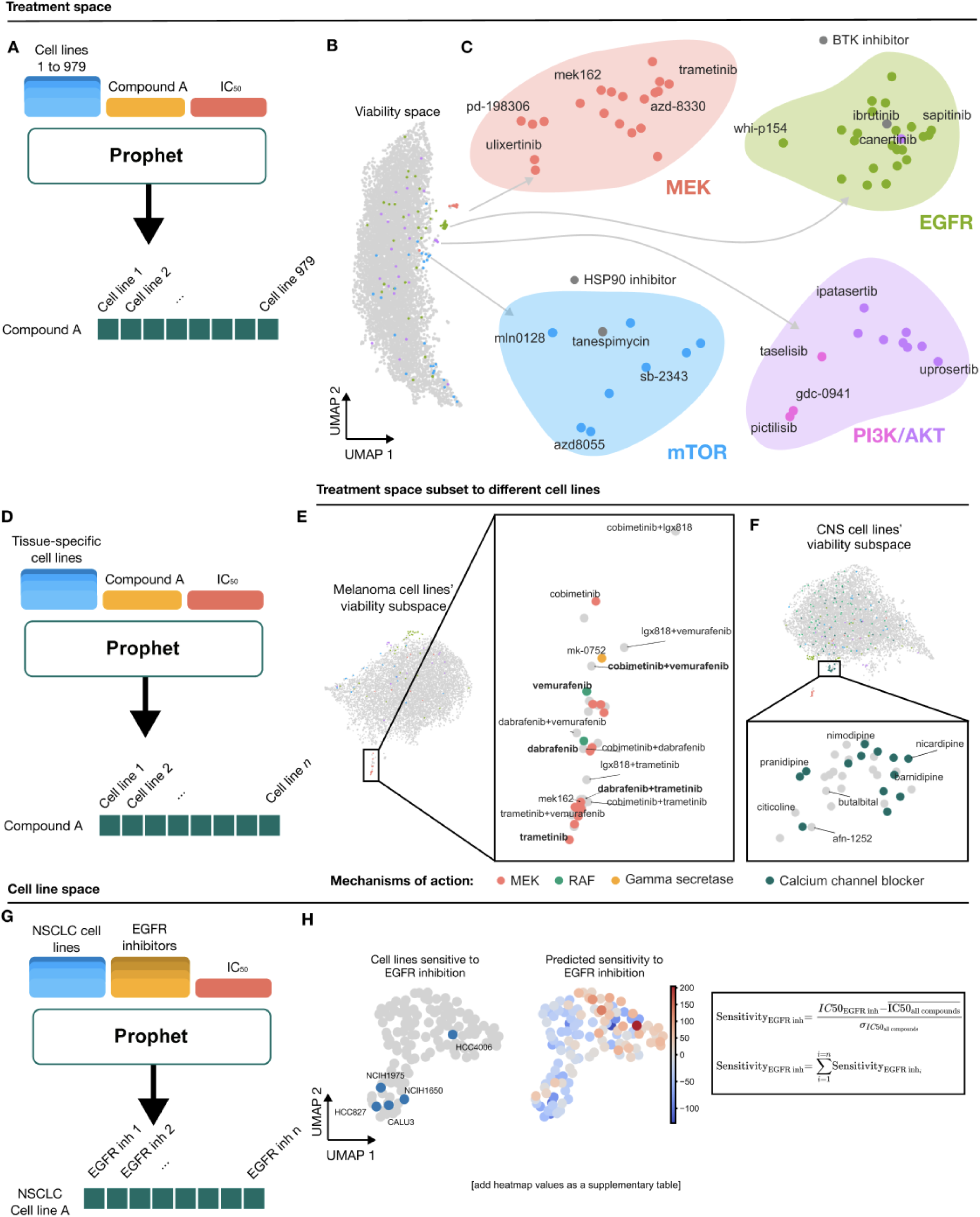
Prophet learns a map of the experimental space. (**A**) Schematic of how Prophet generates a complete experimental space *in silico*. For the perturbation space, Prophet predicts the IC_50_ for each compound or compound combination across 979 cell lines. Each single or combinatorial perturbation is then represented as a 979-dimensional vector, in which each dimension is Prophet’s IC_50_ prediction for that compound in one cell line. We call this perturbation representation an “activity-based fingerprint”. (**B**) Two dimensional visualization (UMAP) of activity-based fingerprints of 8,620 small molecule perturbations, each represented as one point. (**C**) Compounds with differential effects can cluster into four different biological signaling pathways involved in cell proliferation: MAPK, EGFR, PI3K/AKT and mTOR (Table S6). (**D**) Tissue or cell state specific viability spaces can be constructed by subsetting to only the predictions from those cell states For example, for the melanoma and brain cancer subspaces, only cell lines representing the respective tissues are included. (**E**) IC_50_ perturbation subspace for melanoma cell lines, zoomed in on a cluster of compounds containing known cancer perturbations. Each dot is a perturbation represented as a 23-dimensional vector in which each dimension refers to the predicted IC_50_ value in one of the 23 selected melanoma cell lines. Clinically approved monotherapies for *BRAF* V600E mutant melanoma are highlighted in bold. The color of each dot indicates its mechanism of action, as shown in the legend below. (**F**) IC_50_ perturbation space for a subset of brain and central nervous system cancer cell lines, where calcium channel blockers (zoom in) are clustered together and exhibit a differential effect. (**G**) To model cell state space with Prophet, each cell state is represented as a vector in which each dimension is Prophet’s IC_50_ prediction for one compound in that cell line. (**H**) Here we model the activity of EGFR inhibitor perturbations in non-small cell lung cancer (NSCLC) cell lines. Each point is a cell line represented as a vector in which each dimension is the predicted IC_50_ value for each perturbation in that cell line for a set of NSCLC cell lines and EGFR inhibitors. The cell lines HC4006, NCIH1975, NCIH1650, HCC827 and CALU3 are known to be sensitive to EGFR inhibitors. Points are colored by predicted IC_50_ sensitivity (Methods) to EGFR inhibitors. Lower IC_50_ values imply higher sensitivity and higher cell death.

### Compounds targeting cell proliferation

We used Prophet to generate viability predictions for 8,620 small molecule perturbations in 979 cell lines (Fig. 3A). We concatenated the 979 cell line viability predicted values per drug to construct an activity-based fingerprint, which together form a viability-contextualized small molecule manifold (Fig. 3B). We focus on four distinct clusters in this space (Fig. 3C, fig. S8A) containing a total of 64 compounds that target specific regulators of cell proliferation: two clusters from the MAPK signaling cascade (containing compounds targeting MEK and EGFR), and two clusters from the PI3K/AKT/mTOR pathway. Two of the 64 compounds have never been seen in the training data (fig. S8B). Of the 8,620 compounds tested, Prophet was trained on 3,039 compounds, each observed in an average of 365 out of 979 cell lines, and was able to accurately complete 86.9% of the experimental cell line × drug space, grouping compounds in an unbiased manner.

Prophet’s identified clusters are also biologically precise: after annotating the 64 compounds (Table S7, Supplementary Methods, fig. S8A), we found only one potential misclassification: the HSP90 inhibitor tanespimycin, which Prophet identifies as similar in effect to mTOR inhibitors^32^. Additionally, we noted one misannotation: Ibrutinib, typically labeled as a BTK inhibitor, has been shown to bind to EGFR in non-small cell lung cancer (NSCLC) ^33,34^. We suggest that Ibrutinib also has a similar effect in other cancer types. The clustering arises from shared phenotypic effects rather than chemical structure, as the compounds show substantial dissimilarity in chemical space (fig. S8C, Supplementary Methods). This indicates that the experiment representations learned by Prophet are driven by the phenotypic effects of small molecule perturbations rather than their chemical structures, underscoring the utility of this manifold in capturing biologically relevant information which could not be identified using structural similarity alone.

### Therapeutic treatment candidates in BRAF-V600E mutant melanoma

We wondered whether we can identify compounds which phenocopy existing, clinically approved perturbations. To this end, we examined Prophet’s predictions for four compounds already clinically approved for *BRAF* V600E*-*mutant melanoma. From the full 979 cell line viability space above, we create a subspace of 23 melanoma cell lines categorized as either *BRAF* V600E-mutant or -wild-type (Fig. 3E). 74% of these experiments were unseen in training. Such cancers respond well to various MEK and BRAF inhibitors ^35–37^. This subspace reveals a cluster of compounds containing three clinically approved monotherapies ^36,38,39^ and two combination therapies ^37,40^ which are highlighted in bold (Fig. 3E). 45% of the other compounds in this cluster have been indicated for *BRAF* V600E melanoma in *in vitro* experiments (Table S8), suggesting that the remaining compounds are relevant therapeutic candidates.

We further examine the predicted viability per cell line. Two out of 6 wild-type cell lines were on average sensitive to compounds in this cluster, compared to 9 out of 17 *BRAF* V600E-mutants (fig. S10A). Among these, resistance to MEK or RAF inhibition despite the *BRAF* V600E mutation has been documented in certain cell lines, such as A2058’s resistance to MEK inhibitor PD0325901^41^, a behavior our model accurately predicts (fig. S10B) even though the current mechanism of resistance remains unclear.

### Predicting brain and CNS-specific monotherapies

A similar *in silico* generated subspace of perturbation effects in brain and central nervous system (CNS)-derived cell lines reveals a cluster of neurological compounds, enriched for putative calcium channel blockers (p-value<10^−7^, Fig. 3F, Supplementary Methods) which did not cluster when looking at global tissue responses (fig. S9C). Calcium signaling promotes growth and tumorigenesis of brain cancers^42^. Recent studies demonstrate that calcium channel inhibition suppresses brain tumor growth, decreases the expression of oncogenes and increases expression of tumor suppressor genes, even proposing to repurpose existing drugs to treat brain cancers ^43,44^.

### Experimental space contextualized by RNA expression

While the compound space detailed above was constructed using Prophet-predicted viability values, Prophet can also construct a compound space using other phenotypes, such as RNA expression learned from the LINCS dataset. We make predictions for BIRC5 (a gene which encodes the survivin protein) expression after perturbation in the same set of compounds and cell lines (fig. S9D), expanding the original by 5,846 small molecules and 922 cell lines. This new space captures patterns of BIRC5 expression after compound perturbation (fig. S9E). This enables us to identify a set of MEK inhibitors affecting BIRC5 expression and reveals a new cluster of compounds acting on BIRC5 (fig. S9F). Although BIRC5 expression after perturbation has not been reported in the literature for many of these compounds, the cluster includes a survivin inhibitor and Auranofin, which has been reported to downregulate BIRC5 in osteosarcoma^45^. Both of these compounds were not previously tested in LINCS.

### Identifying EGFR sensitivity in non-small cell lung cancer

Applying drug activity fingerprint concept to cell lines, we can also generate a fingerprint for cell lines which we can use to investigate the response signatures of biological systems to a specific array of drugs. As an example, we identify non-small cell lung cancer (NSCLC) cell lines that react to EGFR inhibition. We subset the experimental space to NSCLC cell lines and compute an average EGFR inhibition sensitivity score per cell line (Fig. 3G, Supplementary Methods). *EGFR* mutant cell lines, with a constitutively active EGFR receptor, are often especially sensitive to EGFR inhibitors ^46^. Prophet identifies a gradient in the cell lines with respect to sensitivity to EGFR inhibitors, and is successful at scoring five known putative *EGFR* mutant or EGFR-sensitive cell lines as EGFR sensitive (H1975, H1650, HCC4006, HCC827 and CALU3) ^47–49^(Fig. 3H).

### Virtual screening and in-vitro validation of melanoma-selective compounds

To further demonstrate Prophet’s potential for in silico experimentation and therapeutic discovery, we used it to computationally prioritize compounds for the treatment of melanoma, a highly aggressive skin cancer with tens of thousands of new cases diagnosed annually^50^. By simulating a drug screen in silico, Prophet enabled us to nominate compounds whose effects may replicate or complement clinically approved therapies. We then experimentally validated a subset of these predictions in vitro, showcasing the model’s ability to guide targeted discovery efforts.

To do so, we conducted an in silico screen of all 1.9 million small molecules in ChEMBL (Fig. 4A, B, D), in 1406 cell lines, for a total of 2.7 billion experiments. Only 0.12% of these experiments were contained in the training data. We then selected compounds predicted to have a similar activity fingerprint in melanoma cell lines to existing clinically approved MEK inhibitors (Methods). This resulted in a selection of 123 compounds, none of which had been seen during training (Table S10). However, due to vendor unavailability, not all compounds were purchasable (Table S10). To validate Prophet’s predictions in the respective cell lines, we tested five compounds in HeLa, A375 and SK-MEL-2 cells: CHEMBL3335206^51^, CHEMBL3645238^52^, CHEMBL244488, CHEMBL450775, and CHEMBL507836^53^. With the exception of CHEMBL3335206, which showed no effect on cell viability, CHEMBL3645238, CHEMBL244488, CHEMBL450775, and CHEMBL507836 drastically reduced the viability of A375 and SK-MEL-2, but not HeLa cells (Fig. 4C), mirroring previous reports. Annexin-V staining on CHEMBL507836 and CHEMBL3645238 confirmed that these compounds induce apoptosis, consistent with MEK inhibition (Fig. 4E, F).

**Fig 4.**
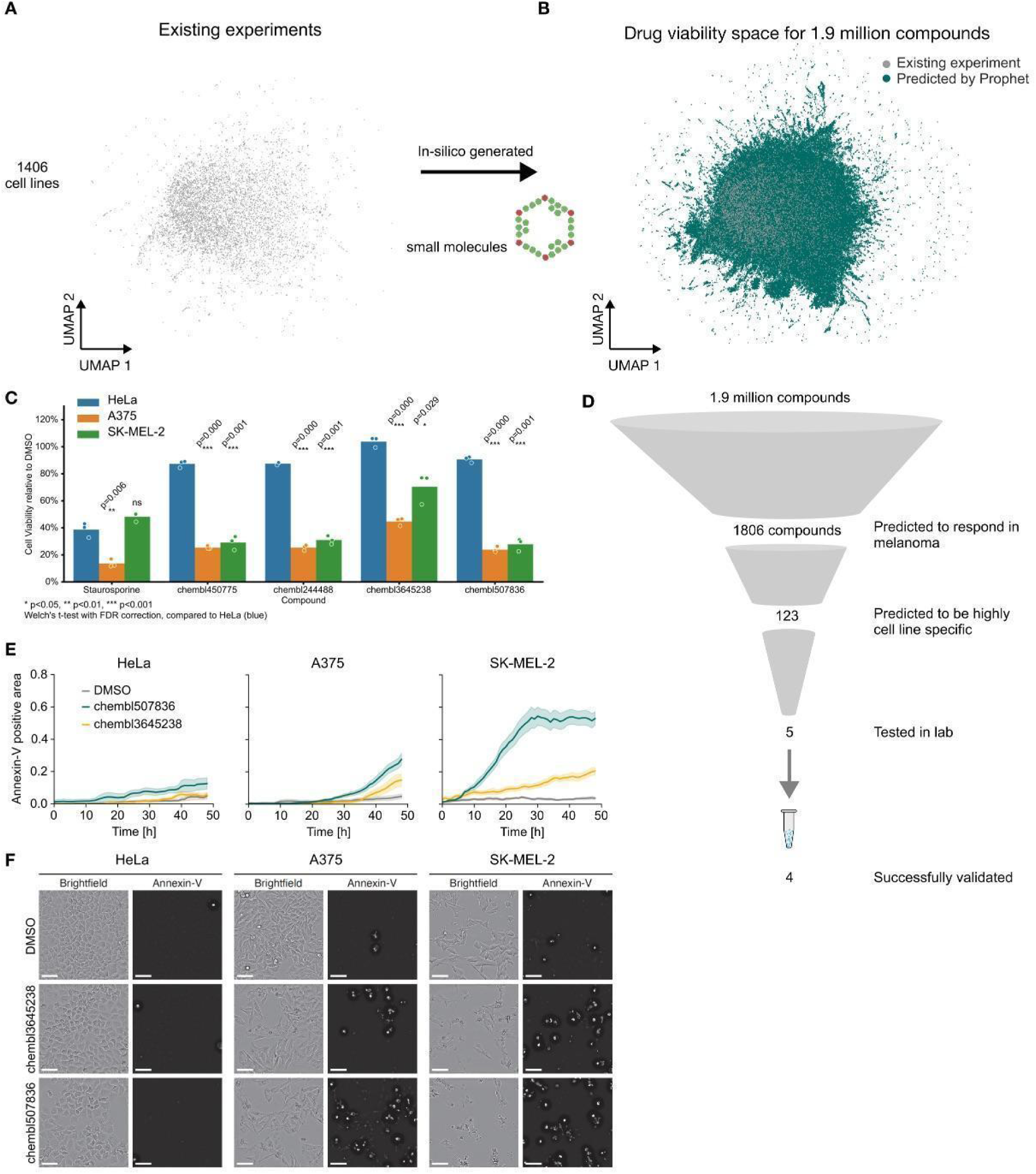
Unseen compounds validate in vitro for melanoma-specific killing. A) UMAP visualization of the viability space across CCLE cell lines of compounds seen in training. Missing cell lines and compound experiments were filled in by Prophet. B) Teal - compounds as predicted by Prophet. Grey - compounds seen in training. The small molecule experiments seen in training cover only 0.12% of the space of possible cell line and singleton drug perturbation experiments possible with 1.9 million ChEMBL small molecules. C) Results from in vitro validation of cell line killing of compounds selected by Prophet and a positive control. D) Prophet’s in silico screen started predicting 1.9M compounds. Among those, 123 were predicted to have a highly specific cell line effect similar to clinically approved therapies. 5 were selected for wet lab validation in melanoma cell lines and 4 successfully validated. E) Annexin-V staining over time in resistant (HeLa) vs. sensitive (A375, SK-MEL-2) cell lines. F) In the same three cell lines as D, microscopy images from two of the compounds with paired Annexin-V staining, demonstrating activation of apoptosis. Scalebars represent 75µm.

These results highlight Prophet’s ability to conduct in-silico experiments within its learned perturbation space, identifying candidate compounds with desirable therapeutic profiles. By modeling perturbation effects across chemical and phenotypic spaces, Prophet enables large-scale in silico screening that can uncover novel therapeutic candidates and plausible mechanisms of action. This positions Prophet as a valuable tool for accelerating data-driven drug discovery.

### Zero-shot in vitro validation under experimental constraints

Building on the previous validation work, we next tested Prophet’s predictions in a more constrained drug screening setup. To do so, we used a setup with existing pre-built 384-well plates, each with 352 unique drug perturbations applied to 9 cancer cell lines (Table S9). In total, there were 16 pre-built plates. A pre-built screening plate contains a library of compounds which have already been fixed onto the plate. As such, compound selection is restricted to predefined sets of 352 compounds instead of 1 compound at a time. This setup represents a scenario in which limitations in experiment size constrain the compounds and cell lines that can be tested.

Aiming to pick a plate with maximally diverse drug responses across cell lines, we used Prophet to make *in silico* predictions for the outcomes of the compounds in each plate in the nine available cell lines. We used the predicted effects on cell viability to score each plate in two categories—dissimilarity across compounds and similarity within clusters (fig. S11A, Supplementary Methods). We then tested the selected plate and compared the results to Prophet’s predictions. Prophet successfully predicted the overall outcome for the entire plate, with a median of 0.53 Spearman correlation (Fig. 5C), despite not having seen any related data during training. Comparing the predicted viability scores for each compound of the selected plate with viability measurements in PRISM, the dataset with the best coverage of the compounds and cell lines out of which 8/9 cell lines and 41/352 compounds were also tested, resulted in a median of 0.07 Spearman correlation (Fig. 5C).

**Fig. 5.**
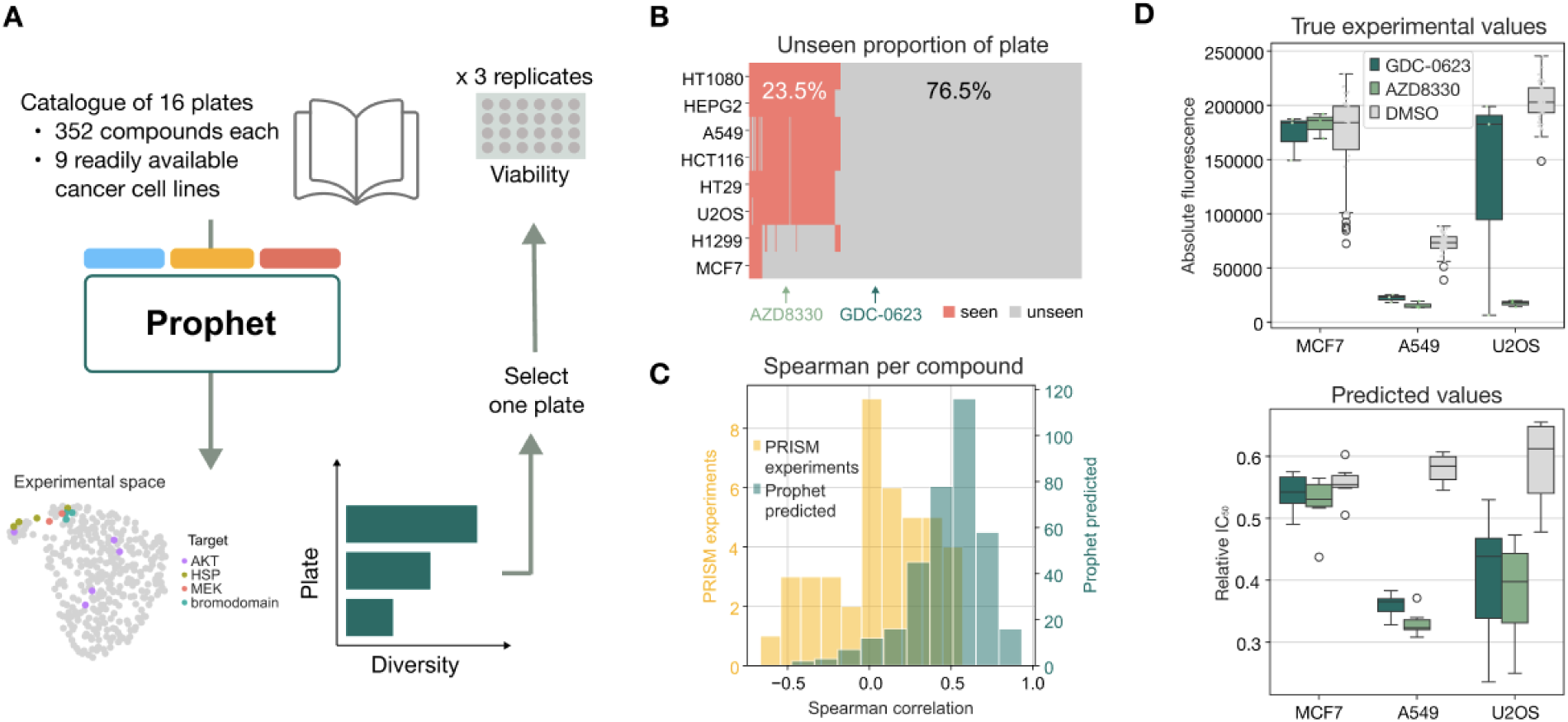
In vitro validation of selected Prophet predictions. (**A**) Workflow accommodating the constraints of an in-house small molecule screening setup. (**B**) Proportion of the performed screen with completely new experiments. Cell line and drug experiments not seen by Prophet in any training dataset shown in gray. Experiments which were measured at least once, with any phenotype, are shown in red. Each column represents one compound. The two compounds also in panel D are highlighted. (**C**) Spearman correlation per compound of the real vs. predicted viability values across cell lines (teal, right axis) for the 352 compounds we experimentally validated. In yellow are the same correlations, but with viability values measured from PRISM, for those compounds which were also tested in PRISM. (**D**) Absolute CellTitreGlo fluorescence values from the in vitro experiments (three replicates) vs. Prophet’s predicted values (6 ensembles). Boxes show the quartiles, and whiskers show 1.5 times the interquartile range from the box. Prophet provides tight uncertainty bounds for these predictions, and more loose uncertainty for the U2OS cell line.

For the two known MEK inhibitors tested, GDC-0623 and AZD8330, Prophet correctly predicted no significant change in cell viability after perturbation specifically in MCF7, but none of the other cell lines, with high certainty (Fig. 5D, fig. S11B). Our experimental validation confirmed this observation (Fig. 5D), showing that both MEK inhibitors produced no significant changes in viability in MCF7 cells, but significant changes in other cell lines. This finding is supported by previous work documenting MCF7 resistance to some MEK inhibitors, including trametinib^54^, but not other MEK inhibitors such as selumetinib and pimasertib^55^. Prophet independently uncovers this highly specific resistance despite not previously seeing either compound in MCF7 cells in training, and not seeing GDC-0623 at all.

### Prophet predicts in vivo phenotypes

Prophet’s capabilities extend to *in vivo* experimental settings. Saunders et al. 2023^25^ generated a single-cell transcriptomic atlas of the effect of 23 genetic knockouts—including combinations—on 1,812 developing zebrafish embryos across 5 timepoints (Fig. 6A). Their analysis includes using the single-cell transcriptomics data to compute relative cell type frequencies for each embryo (Fig. 6B) and comparing the genetic perturbations using similarities and differences in cell type proportion change. In this high-dimensional space, genetic perturbations clustered together based on the affected zebrafish developmental lineage. Given our framework, we asked if Prophet could learn these cell type proportion changes and make predictions about additional knockouts for specific lineages.

**Fig. 6.**
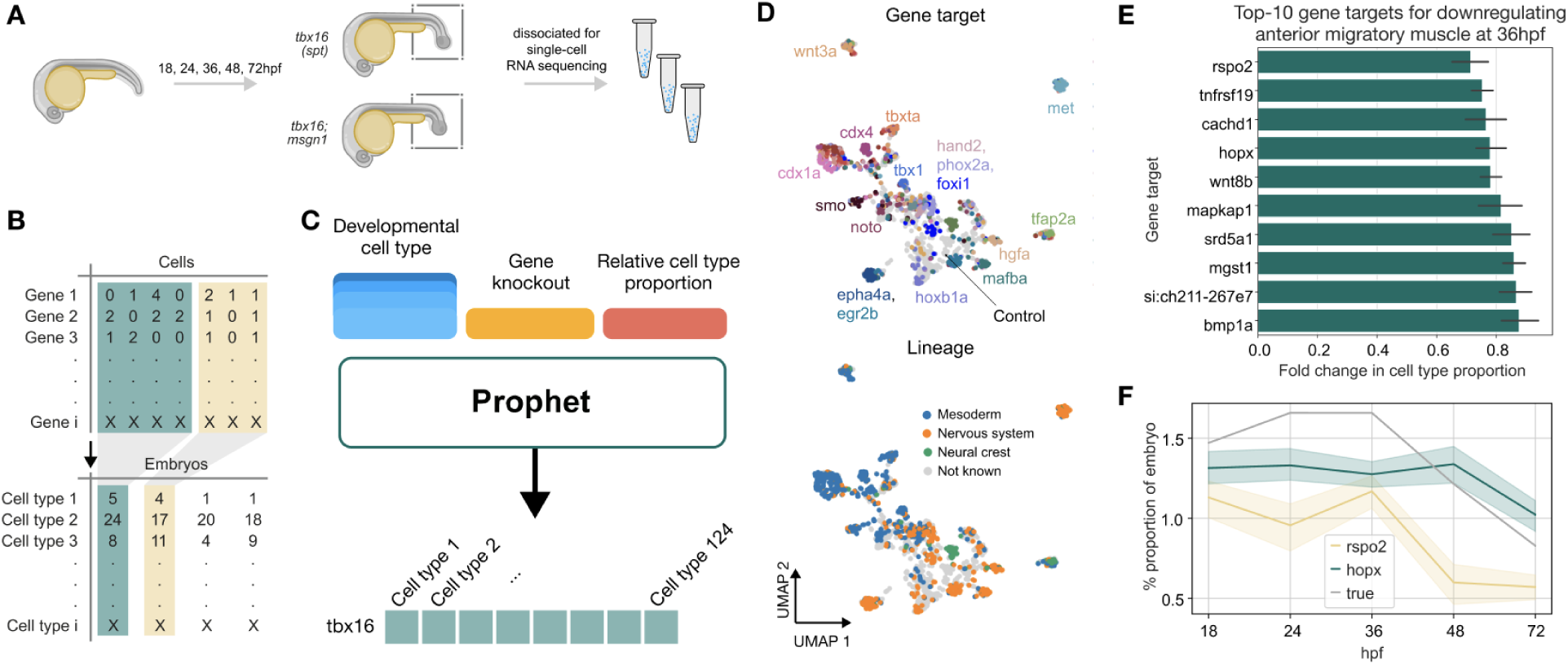
Prophet extrapolates relative cell type abundance in zebrafish development. (**A**) Summary of the original experimental setup from Saunders et al. 2023. (**B**) How the single-cell x gene matrix was converted into cell type proportions per embryo. The latter is the phenotypic value predicted by Prophet. Adapted from Saunders et al. 2023. (**C**) Schematic of how Prophet generates a complete experimental space *in silico*. Prophet predicts the cell type frequencies of one zebrafish embryo for each genetic KO combination across 124 cell types to construct a genetic KO “fingerprint”. (**D**) Prophet’s learnt experimental space of genetic KO “fingerprints”.. Depicted are: combinatorial predictions which include a knockout seen in the original publication (top panel, colored by the seen knockout; bottom panel, colored by the lineage of the seen knockout) and singleton predictions for zebrafish transcription factors (gray) not seen in the original publication. Prophet’s experimental space recapitulates the findings obtained by the authors in the original publication and the separation between genes involved in different developmental lineages. (**E**) Top-10 gene targets that suppress the appearance of the “anterior migratory muscle” cell type as predicted by Prophet, ranked by their average effect on the fold-change of the cell type proportion as compared to control. Rspo2 and Hopx are known regulators of mesoderm development, while Wnt8b and Bmp1a belong to families of genes also involved in the same lineage. (**F**) The predicted absolute cell type proportion of anterior migratory muscle at 5 timepoints for the top 1 and 3 predicted gene targets, with control as comparison.

Given that this is the only *in vivo* dataset and that no common prior knowledge exists for it within our dataset compendium, we did not use any pretrained Prophet version, but trained a single Prophet model from scratch (Methods). We then used the predicted relative cell type frequencies to construct a zebrafish cell type proportion experimental space, extending it with predictions of single knockouts, and analyzed perturbation-dependent deviations relative to wild-type embryos (Fig. 6C).

Despite only 23 perturbations in the dataset, the cell type proportion experimental space learnt by Prophet qualitatively matches the one obtained by the authors in Fig. 2C of the original publication (Fig. 6D, top panel), where it can be seen that different gene knockouts provoke different responses in terms of cell type proportion in the zebrafish embryos. The gene knockout fingerprints learnt by Prophet group genes according to their involvement in the mesoderm, nervous system and neural crest lineages (Fig. 6D, bottom panel). We also run predictions for singleton knock outs of zebrafish transcription factors (TFs): for example, znf711 and znf740b from the zf-C2H2 family and dbpb, bach1b from the bZIP family, to produce additional candidates for further experiments.

As an example, we run Prophet to get predictions of gene targets that reduce the cell type proportion of anterior migratory muscle at 36 hours post-fertilization (hpf)(Supplementary Methods). We measure the strength of a knockout as fold change using the control embryo cell type proportion at 36hpf as a reference. Among the top 10 gene targets predicted (Fig. 6E), we find that Rsp02 and Hopx rank first and third, respectively; for both genes, previous studies describe their key role in mesoderm development^56,57,58^. Other genes in the top-5 (Bmp1 and Wnt8b) are part of families of genes also involved in mesoderm development^59–61^. All ten of the Prophet-predicted strongest gene knockouts are not part of the original dataset. Prophet’s predictions also report an uncertainty obtained from the variance of predictions of an ensemble of models (Methods). This informs experimentalists of Prophet’s confidence in a prediction, allowing them to make an informed decision about whether or not to include high uncertainty TFs in a panel.

Finally, Prophet can also provide fine-grained predictions across timepoints by incorporating the time of the phenotype into the phenotype token. The proportion of anterior migratory muscle cells varies across time points given an F0 knockout in a zebrafish embryo in the genes Rspo2 and Hopx (Fig. 6F). These detailed temporal predictions can assist biologists to make more informed decisions about which subsequent experiments to conduct, such as focusing on particular stages for gene expression analysis or investigating downstream effects during critical developmental windows.

This example showcases how Prophet is flexible enough to extend beyond cell lines and *in vitro* experiments to *in vivo* phenotypes. In the future, Prophet could be provided with genetically orthologous data in human developmental models to bridge the gap between controlled laboratory experiments and complex *in vivo* systems, helping to translate findings from basic research into humans and potential therapeutic applications.

## Discussion

Recent advances in foundation models have transformed fields like natural language processing^15,16^, vision^62^, protein design^63^ and molecular cell biology^53,64^, largely driven by transfer learning from large, diverse datasets. Inspired by this paradigm, Prophet introduces a novel and scalable approach for phenotypic experimental biology, leveraging transfer learning to effectively predict outcomes across a wide range of biological experiments. Pretraining on millions of experiments and fine-tuning for specific datasets allows Prophet to consistently outperform simpler architectures and mean baselines on unseen conditions (Fig. 2E, Fig. S3). This enables large-scale in silico experimentation: for example, Prophet was used to screen 1.9 million compounds from ChEMBL across 1,406 cell lines, simulating 2.7 billion experiments, to nominate candidate MEK inhibitors for melanoma. Experimental validation confirmed four out of five tested compounds selectively reduced viability in melanoma cell lines, demonstrating the practical utility of Prophet’s predictions in guiding targeted discovery. Such capabilities also allow smaller in silico screens to recover the same number of hits up to 60× more efficiently (Fig. 2G, fig. S5B), while activity-based compound fingerprints accurately recover known biological pathways. Notably, Prophet extends beyond cell lines to in vivo settings (Fig. 6), demonstrating the potential to systematically map complex experimental landscapes.

Limitations of the current approach include the simplified assumption of the three-axis experimental space (Fig. 1). Other elements, such as temporal dynamics, dosage, or technical variations, are also part of each experiment. Currently, Prophet treats time points as part of the phenotype and does not consider their ordinal nature; dosage could be treated similarly in future extensions. Prophet could also incorporate raw genetic sequences or chemical structures, extend to antibody perturbations, or explicitly account for technical covariates such as incubation period or assay conditions. Presently, Prophet accommodates technical differences by assigning distinct phenotype tokens per dataset, allowing the model to learn dataset-specific effects (e.g., IC50 measured via CellTitreGlo in GDSC2 versus Luminex fluorescence in PRISM). Prophet is not designed to correct for intra-study experimental noise, assuming that data preprocessing and quality control have already been applied. Additionally, it does not currently implement formal Bayesian uncertainty estimation; ensemble-based approximations of predictive uncertainty provide only a rough estimate, with correlated members. These limitations represent opportunities for future development, particularly as more perturbational data becomes available, enabling iterative improvements of Prophet’s mapping of experimental landscapes.

While Prophet successfully transfers learning between independent assays and phenotypes, a major challenge remains in transferring knowledge across scales—between in vitro, in vivo, and patient-derived experiments. Mastering this transfer is crucial for clinical translation, as complex functional assays (organoids, patient samples, in vivo models) are often limited in size. Prophet’s architecture is well-suited to leverage large-scale in vitro perturbation screens to inform predictions in these more complex biological contexts, enabling more accurate and generalizable modeling of experimental outcomes.

Prophet establishes a new paradigm in phenotypic drug discovery. By integrating diverse datasets under a single learning framework, it enables large-scale virtual screens, activity-based compound fingerprinting, and targeted experimental design. Experimental validation demonstrates that Prophet can prospectively identify therapeutically relevant compounds, such as MEK inhibitors for melanoma, confirming both its predictive accuracy and utility for lead discovery. As a flexible backbone for lab-in-the-loop systems, Prophet can guide the next generation of data-driven experiments, prioritize efforts, and de-risk experimental campaigns, paving the way for systematic exploration of biological perturbation space.

## Acknowledgments

Madeleine Duran and Cole Trapnell for advice on zebrafish analysis and the list of annotated zebrafish transcription factors. Paul Bertin for suggesting the use of an accuracy score for biological interpretability, additional potential model architectures, and innumerable other helpful insights. Szen Toh for making suggestions for both datasets and embeddings, helping coordinate with the Garnet lab which authored so many of these datasets, and commenting on the initial draft of the manuscript. Tim Treis for invaluable advice on the cell painting datasets. Luke Zappia for helpful and extensive commentary on the figure designs. Stefanie Brandner for excellent technical assistance in the small molecule library experiment. Riccardo Trozzo, Ekaterina Zhigalova, and Juan José Montero Valderrama for biological advice on CRISPR perturbations. Stefan Peidli, Tessa Green, and Camille Fourneaux for advising on content and clarity in the initial draft of the manuscript.

## Author contributions

F.J.T. conceived the study together with A.T.L. and Y.J.; A.T.L. and Y.J. contributed equally and have the right to list their name first in their curriculum vitae; F.J.T. supervised the project and acquired funding; A.T.L., Y.J., F.J.T., and S.K. developed the methodology; A.T.L. and Y.J wrote the whole code; S.K., Y.J., A.T.L., I.R., J.T., and K.H. carried out the investigation; Y.J., A.T.L., X.Z., and Z.Z. performed the visualizations; N.S. performed the in vitro validation experiment; Y.J., A.T.L., and N.S. wrote the original draft. All authors read, corrected, and approved the manuscript.

## Data and materials availability

The code for Prophet can be found at https://github.com/theislab/prophet. Data obtained from in vitro validation can be found at https://figshare.com/articles/dataset/Prophet_viability_plate_xlsx/26516002

## Methods

### External datasets

We selected datasets with at least 100 clearly labeled cell states or perturbations. In most cases, we used as the predictand the least processed values post-batch correction generated by the original authors. The contents of each dataset are detailed in Table S1.

All datasets then underwent the following: replicates were averaged and experimental outcome values were scaled from 0 to 1. Samples with a cell line, perturbation, or phenotypes which we did not have an embedding for were filtered out. For example, small molecules without SMILES strings or with multiple SMILES strings were filtered out.

### Baseline models

The baseline models we employ are linear regression, a multilayer perceptron (MLP) and Random Forest. As some experiments are combinations of treatments and the baselines are not permutation invariant, all datasets are duplicated and the interventreatmentstions flipped. This means that an observation 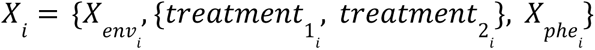 is also seen as 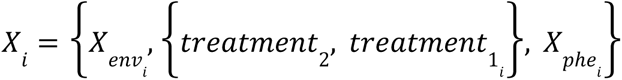. For the linear regression setting, L1 and L2 regularization are tested but just the best results are reported. Random Forest runs with 100 estimators, minimum samples per split 2 and without capping the max length.

### Data splits

We distinguish between two distinct stratifications: unseen perturbations and unseen cell states. For unseen perturbations, the test dataset consists entirely of new perturbations - genetic perturbations or small molecules. In combinatorial datasets such as zebrafish, any combination including the unseen perturbation is placed in the test set. For unseen cell states, the test dataset consists of experiments in entirely new cell states.

In both cases, the test dataset represents 20% of the total perturbations or cell lines, while the validation dataset encompasses 10%. The only exception is the zebrafish embryo dataset, where a single gene was held out at a time instead of 20%. For instance, when *tbx16* was the held-out gene, the test set contained all conditions where *tbx16* was knocked out: *tbx16*, *tbx16+tbx16l*, and *tbx16+msgn1* crispant embryos. This ensures that evaluation is conducted on truly unseen gene targets, with validation remaining at 10% of genes.

When fine-tuning a pretrained Prophet model, we ensure no data leakage or confounding issues by maintaining consistent train and test splits between the large pretrained model and the assay-specific models.

Additionally, we also assess the model’s performance as it is exposed to progressively more perturbations or cell lines (fig. S4,5), evaluating Prophet with 80%, 50%, 30%, 20%, 10%, and 5% of the perturbations or cell lines included in the training dataset.

To further assess the generalization capabilities of Prophet, we devised 5 additional stratifications based on what would typically be considered a difficult task.

- Unseen scaffolds - Each small molecule is assigned a Murcko scaffold, which captures its core structural framework. Drugs in the test set have Murcko scaffolds that are not present in the training set, ensuring structural novelty. This stratification strategy is only applied to small molecule datasets.
- Unseen tissues - Whole CCLE lineages (e.g. skin, stomach, liver) are excluded from training and used as test. Only applied to datasets with at least 5 cell lines.
- Unseen drug clusters - each small molecule is assigned a drug cluster based on the leiden community membership of its chemical structure embedding used by Prophet. This leaves out a chunk of input space, making prediction especially hard for the model.
- Unseen gene clusters - a similar concept as unseen drug clusters, but using genetic embeddings.
- Unseen cell line clusters - a similar concept to unseen drug and gene clusters, but using cell line embeddings. Differs from unseen tissues where cell lines did not cluster by tissue of origin.
- Both cell line and perturbation are unseen - similar to unseen perturbations, except the cell lines are also unseen.

### Performance evaluation

We use three primary metrics in evaluating Prophet’s performance: hit rate, R2, and Spearman correlation.

Of these, hit rate is the most relevant and interpretable. For each test split, we took the top 5% of the unseen covariate (e.g. for unseen compounds, the compounds which produced the top 5% of values per cell line) and scored how many of those Prophet identified as being in the top 5% within its predicted top 5% (e.g. how many of the unseen compounds Prophet also ranked in the top 5% for that cell line). This provides us with a new metric we call “hit rate” which is related to both precision and recall (mathematical derivation below). We then did the same for the bottom 5%, and averaged the hit ratio between top and bottom. For example, when asked to predict the IC50 of 100 unseen compounds in A549, the top 5 and bottom 5 compounds according to IC50 are considered hits. We use Prophet to predict IC50 values for the 100 compounds, and then take the top 5 and bottom 5 predicted compounds as Prophet predicted hits. Prophet predicts 3 of the top 5 compounds within its top 5 compounds, getting a hit ratio of .6. Prophet predicts 2 of the bottom 5 compounds within its bottom 5 compounds, getting a hit ratio of .4. This is averaged into a hit ratio of .5, and then the mean (.05) is subtracted to get a value in the table of .45. This is performed, per dataset, for every cell line and every phenotype in every test split to get an overall averaged hit ratio. This means, for instance, that in the case of JUMP, in which we predict the Cell Painting features, we predict the individual Cell Painting features, compute the hit ratio for every cell line and every feature, and then average its results, resulting a single number.

Formally, let:

- 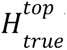 be the set of *topk*% true hits,
- 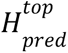 be the set of *topk*% predicted hits,
- 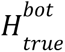 and 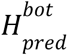 be the analogous sets for the *bottomk*% of values.

We compute:

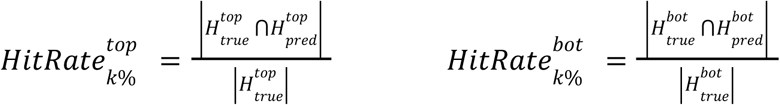

The overall hit rate is the average:

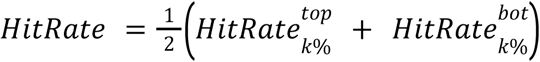

Overview of Prophet

Prophet considers a perturbational dataset of *N* experiments 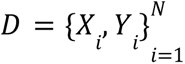 where *X_i_* represents the experiment carried out and *Y*_*i*_∈*R*^*K*^ is the k-dimensional observed phenotype. Each experiment *X_i_* is modeled as a set 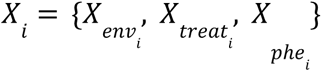 conformed by the environment where the experiment is carried, on 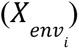, the set perturbations applied 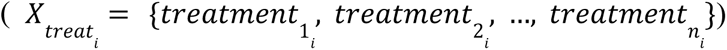 and the phenotype that is measured 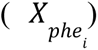. The goal of Prophet is to learn a function that maps a novel experiment *X_i_* to an observed phenotype *Y_i_*.

Given an experiment description 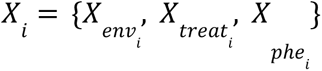, Prophet first projects all representations into a common token space *Z*_*i*_∈*R*^*d*^, where each token represents an element of the experiment, using the tokenizer networks. The perturbation and cell state typically have prior representations, which Prophet transforms using feed-forward networks. In contrast, the phenotype does not usually have a predefined representation. Instead, Prophet models it as a learnable embedding. Hence, 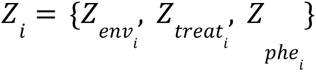 where 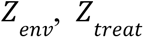 and 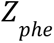 are the outcome of the tokenizer networks. In the case of the set of treatments applied *Z* _*treat*_, <PAD> tokens are appended if needed to ensure a consistent length in all observations. Subsequently, we prepend a learnable class token <CLS> into the sequence of experiment-related tokens, resulting in the augmented set 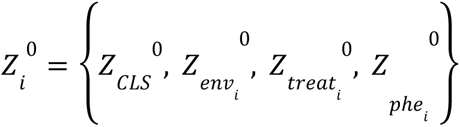 The set 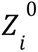 is then fed into the transformer, where the combination of different feed-forward networks, residual connections and self-attention modules produce an output representation 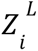 at the final layer *L* of the transformer.

The resulting final state of the learnable <CLS> token 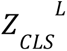 serves as a comprehensive representation of the experiment. Alternatively, we also tried linear addition of all output tokens instead of using only the final state of the <CLS> token, which led to very similar results. This representation is employed for predicting the observed phenotype *Y*­*_i_* through a function *f*: *R^d^* → *R^k^*. The entire network is trained end-to-end to minimize the MSE error between the predicted experimental outcome *Ŷ_i_* and the real experimental outcome *Y_i_*.

Prophet does not use any positional encoding. Instead, it models the experiment as a set of elements without any specific intrinsic order (environment, treatment, phenotype). Self-attention is a permutation equivariant operation, so altering the order of the elements in the set does not alter the final result.

### Tokenization

We call tokenization the process of projecting elements of the set 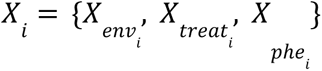, that represents an experiment, into a common token space to get representations *Z_i_* ∈ *R*^*d*^.

For each of these elements, a specific tokenizing function *f*: *R*^*p*^ → *R*^*d*^is learnt where *p* is the dimensionality of the initial representation and d is the dimensionality of the token space. Those functions project each element of the experimental set into the common token space that serves as input for the Transformer. In the case of the treatments, they undergo a different transformation *f* depending on whether it is a small molecule or genetic perturbation. All the transformations are modeled as feed-forward networks.

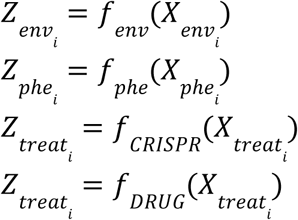

### Cell state representations

The most important determining factor in choosing cell state representations is the size and therefore ability to allow for unseen cell state predictions. For example, choosing a representation of cancer cell lines which only spans 100 cell lines would limit predictions to those 100 cell lines. As such, we used bulk RNA-seq from the Cancer Cell Line Encyclopedia (CCLE) ^656565^ from Dependency Map, which covers 1406 cell lines. To reduce training time, we reduced the dimensionality of both sources using PCA to 300 components. Further details can be found in Table S1.

For the one perturbation dataset performed in zebrafish embryos, we consider each cell type as a different tissue. To do so, we used the pseudo-bulked expression of the cell type in a control embryo across all time points as the cell state representation.

### Treatment representations

All assays examined in this study involve perturbations induced by either CRISPR or small molecules. Hence, the prior representations of each class of perturbations are different to gather the relevant aspect of each. In both cases, however, the different representations are concatenated.

For CRISPR treatments (including KO, interference, and activation), we use the target gene to represent the treatment. We further go on to use two orthogonal sources of information to represent a target gene - its co-expression patterns across cell lines, and its coding sequence. Co-expression pattern representation is derived from CCLE. The coding sequence representation is derived from Enformer embeddings for the whole transcription sequence. Enformer embeddings are derived from the second to last layer of the model. The genomic coordinates of GRCh38 (hg38) were extracted using the BioMart Data Mining tool. In this process, we selected the Human genes GRCh38.p14 dataset and only included genes with an HGNC symbol ID. For genes with multiple transcripts, we retained the coordinate with the smallest Transcription Start Site (TSS). Subsequently, we retrieved the corresponding gene sequences from the GRCh38 human reference sequence produced by the Genome Reference Consortium.

For small molecules, we use the canonical SMILES string to represent the treatment. We further go on to use different representations of molecular structure - RDKit2D<u>(*29*)</u> molecular graph embeddings and Morgan fingerprint embeddings(*30*). RDKit embeddings produced a 200-dimensional vector, while the Morgan fingerprints were calculated as 1024-dimensional vectors with radius fixed at 3. Specifically, we use 195 dimensions that represent the RDKit2D fingerprints and 1024 dimensions for the Morgan fingerprints. In cases in which two datasets used the same common name for a compound but attached different SMILES, we appended a suffix to the compound name (e.g. imatinib_GDSC and imatinib_PRISM) to differentiate between the two.

### Phenotype representations

Phenotypes are typically learned from scratch without any prior knowledge-based representation, primarily due to the challenge of identifying a meaningful prior that accurately reflects the phenotype. Hence, phenotype representations are modeled as learnable embeddings and developed during training.

### In vivo phenotype prediction

In the case of the cell type proportion in the zebrafish dataset, we do not use the pretrained Prophet version since it is a completely different organism than the ones in the pretraining corpus data and the data representations fundamentally change.

In this case, we use a timepoint representation which is the vector of cell type proportions of the control embryos at that timepoint. The cell state embedding is constructed by subsetting to control embryos, computing the top 300 highly variable genes, and then taking the pseudobulk RNA expression of the cell type. Pseudobulks were then normalized with sc.pp.normalize_total with default parameters. The treatment embedding is constructed from the 300 component PCA of the transposed cell x gene matrix of the control embryos, exactly as was done for the human CRISPR treatment embedding. Finally, each readout is the cell type proportion after a given F0 knockout at a given time *t*. To ensure no data leakage in evaluation, the cell type proportions of the control embryos are never used in test or validation.

### Model training

Prophet is efficiently trained in a single Nvidia A100 40GB GPU. We train the model using float32 precision. We use the AdamW optimizer^66^ with β_1_ = 0. 9 and β_2_ = 0. 999, weight decay of 0.01, and dropout of 0.1. The batch size is 4096 in the case of the model training across all assays, while it depends on the size of the dataset in the case of the smaller experiments. The learning rate increases linearly until 1e-4 with a linear warmup of 10,000 steps. After the warmup, a cosine decay regime^67^ is applied. All weights are initialized using Xavier initialization^68^ with default parameters, while the bias terms are initialized to 0. It is run with an early stopping with patience of 20 epochs. During pre-training, it tracks the average R2 performance in the validation set per assay. However, we have noted that model selection after a sufficient number of pre-training epochs does not change results significantly. For fine-tuning, the early stopping tracks the dataset-specific R2 in the validation set, the same early-stopping procedure used in the Prophet-individual models. During fine-tuning, all parameters of Prophet are updated for 20,000 steps with a batch size of 256 observations. These same settings were used for each dataset.

The pre-trained Prophet model consists of 8 transformer encoder units with 8 attention heads per layer and a feed-forward network size of 1024 to generate a 512-dimensional embedding of the experiment. However, the Prophet models trained from scratch in specific datasets are much smaller (2-4 layers and 1-4 attention heads). This is because using a large model in dataset-specific cases resulted in significant overfitting (Supplementary Methods). We also observed that scaling beyond ∼20M parameters in the pre-trained setting offered no performance benefit (fig. S12). This, taking into account the previously mentioned overfitting of the large model in the dataset-specific case, suggests that model capacity is currently constrained by data availability rather than architectural limitations.

### Uncertainty

With the exception of zebrafish development, we compute an uncertainty value for Prophet’s predictions by training six individual models on all datasets. Each model maintains the same test set, but the validation set (and therefore the training set) is selected using an arbitrary seed. As Prophet generally exhibits very stable behavior when learning representations robustly represented in the data, the variance of these predictions can be used as a measure of the uncertainty of the model. Alternatively, we also explored Monte Carlo Dropout^69^ as an alternative uncertainty estimation method. However, in our case, it did not produce uncertainty estimates that were sufficiently informative or well-calibrated, and thus it was not included.

We take a similar approach to computing uncertainty in zebrafish development, but because the zebrafish dataset has no overlap with any existing dataset, we compute uncertainty using different test and validation genes with a model trained on only this dataset.

### In silico screening compound selection

Building on the IC50 perturbation subspace described in the previous section, we used Prophet to expand beyond experimentally tested compounds (Fig. 4A) to a broader space encompassing 1.9 million molecules from ChEMBL (Fig. 4B), evaluated across melanoma and non-cancerous cell lines. That subspace contains a representation of small molecules regarding their effect on melanoma and control cell lines, specifically. To start filtering the 1.9 million compounds to a feasible number of candidates to analyse, we decided to identify regions of the subspace capturing known therapeutic activity and focus on those. DBScan is used to find clusters within the perturbation subspace. Of these clusters, two of them were closely correlated and contained the four compounds involved in clinically approved perturbations for V600E BRAF-mutant melanoma: cobimetinib, trametinib, vemurafenib, and dabrafenib. We computed the centroid of those clusters and measured the correlation of each compound in the cluster with the centroid, retaining those more correlated than the mean. This resulted in 1,806 candidate compounds, a tractable number for subsequent analysis.

We then used an ensemble of Prophet models to make predictions for the 1806 compounds and the original 8620 compounds in Figure 3B. We filtered out compounds for which Prophet predicted inactivity or non cell line-specific behaviour. We then used pertpy^70^ to annotate mechanism of action, resulting in 16 compounds labeled as MEK inhibitors. We filtered again by compounds with greater than .75 Pearson correlation with the 16 MEK inhibitors, resulting in 123 compounds. These compounds had an average rank of 13996 (median 362) when using tanimoto similarity with the known MEK inhibitors, and 22 (median 25) when ranked using Pearson correlation of the Prophet-predicted IC50 with known MEK inhibitors.

## Supplementary Methods

### Prophet-individual training

Each Prophet-individual model has been optimized after an extensive hyperparameter search. They have from 2 to 4 transformer blocks and from 1 to 4 attention heads, depending on the specific dataset (Supplementary Table 2). The final models are selected according to an early stopping selection criteria that tracks the R2 score in the validation set, with patience of 20 epochs.

### Comparison of Prophet-individual training

The single-dataset models were trained using the same data splitting and stratifications as full model fine-tuning, maintaining consistent train/val/test partitions. However, while single-dataset models had hyperparameters adjusted per dataset, the fine-tuned Prophet models did not use dataset-specific hyperparameters; all datasets shared the same fine-tuning settings, including 20,000 training steps and a batch size of 256 (reiterated from Methods). Fine-tuning decisions were independent of Prophet-individual results, with early stopping based on dataset-specific validation R² during fine tuning.

### Cell line embedding

We obtain the transcriptomic variability vector by transposing the CCLE “cell line x gene expression” matrix and finding a low dimensional representation of the genes using PCA. We derive the genomic sequence vector from Enformer’s embeddings ^71^ for the entire coding sequence of the gene.

### Alternative cell line and treatment embeddings

For the perturbation embeddings, different prior-knowledge based options were tested, such as generating embeddings from STRING^72^, Perturb-Seq datasets ^73,74^, gene2vec^75^, GO pathways ^76,77^ or transposing the CELLxGENE^78^ matrix and taking PCA to obtain a low dimensional gene representation^79^.

Regarding the cell lines, the copy number variant information provided by the Cell Model Passports^78,79^ was also tested.

None of those options provided better results than the presented settings.

### Quantifying improved experimental design and AUPRC using in silico predictions

We define a logFC hit to be one for which the log fold-change in the phenotype of interest increases or decreases by 2x as compared to DMSO in the same cell line. For example, if A549 has an IC_50_ of .9 to DMSO, only compounds predicted to have an IC_50_ of .45 or lower would be active. We compute the area under the precision-recall curve using sklearn’s ‘PrecisionRecallDisplay’ with the true binary indicator as the 2x threshold computed on real values, and the *in silico* predicted fold-change values as the prediction vector, resulting in a 12.7x increase in the number of logFC hits discovered (recall). To evaluate Prophet’s ability to reduce the number of experiments needed (precision), we used a 4x LFC cutoff. This translated into a 60x reduction in the number of necessary experiments by assuming an experiment in which the user assays the small percentage of compounds predicted to be positives by Prophet, and compares the number of logFC hits obtained to an assay of the same size but in which random compounds from GDSC were chosen (fig. S5B).

### Constructing activity fingerprints

To construct a small molecule viability embedding space, we use the predicted IC_50_ values of 979 cell lines as the vector representation for each compound. Cell lines were selected from those seen at least once in any dataset, and compounds were excluded if we *only* had an InChi key hash or Broad ID, such that we could not biologically interpret their use or activity, nor could we order them for further experiments. We then co-opted the default scanpy^80^ workflow to normalize predicted values per cell line, compute PCA, neighbors (Euclidean distance with 15 neighbors), and finally UMAP^81^.

For the embedding space constructed from chemical structure alone, the RDKit and Morgan fingerprint embeddings used to train the model were used as the vector representation for each compound. We subset the 156k available embeddings to the same 8620 compounds which are also displayed in Fig. 3b and preprocessed them in the same way.

While one could examine the experimental space learned by single-assay Prophet models, exploring the space of the Prophet model trained across a diverse array of experiments offers additional advantages. The larger model benefits from being trained on a more comprehensive dataset (Fig. 2), which enables it to capture a broader range of experimental conditions, leading to more robust and reliable predictions. Hence, its experimental space is better positioned to provide nuanced and contextually accurate insights, making it a more powerful tool for guiding biological research and experimental design. On top of that, it is a single model able to span different phenotype-specific experimental spaces, instead of multiple models creating multiple spaces.

### Metadata annotation of small molecules and cell lines

Compounds were annotated with putative mechanisms of action and protein targets from DepMap^70,82^ using their common name. Common names were retrieved from the respective metadata for each dataset. The following compounds which could not be automatically annotated and for which the name of the compound did not make the mechanism of action abundantly obvious were manually annotated in Supplementary Table 7.

### Enriched mechanism of action annotation

DBscan^83^ is an unsupervised community detection algorithm that identifies clusters according to the density, and importantly does not assign all points to a community. We use the pertpy (Heumos et al. 2024) implementation of DBscan to identify clusters using the following parameters: min_sample = 5, eps = 0.25. The outlier cluster and the largest cluster, representing non-effective drugs, were excluded from the analysis. Clusters in which at least 33% of compounds are annotated with the same mechanism of action or protein target are considered enriched for that pathway. P-values for this enrichment were determined by Fisher exact test.

### Computing cell line sensitivity to a set of compounds

A cell line’s sensitivity to a single compound is defined as

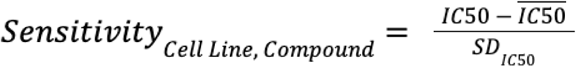

Where IC50 and SDIC50 are the mean and standard deviation of the IC50 values across all 8620 compounds. We assume that most compounds will not have an effect on the cell line.

A cell line’s sensitivity to a class of compounds is therefore:

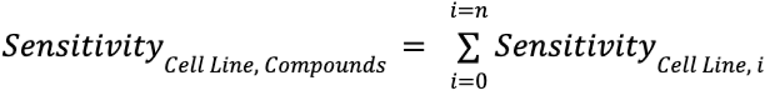

### Scoring sets of 352 compounds for diversity

As nine cell lines were prepared to be treated in triplicate in 384-well plates at 5 µM concentration and incubated for 72hrs (Methods, Table S9), we used Prophet to predict the *in silico* outcome of each of these plates and scored them for diversity. We proceeded with the following criteria after generating in silico predictions for each plate and cell line: First, we scored each plate for how negatively correlated compounds were amongst themselves. Second, we used leiden to assign compounds to communities based on their behavior across the nine cell lines and then took the average of correlations within a community. From these, we prioritized a plate with low correlation across compounds but high correlation within communities in order to optimize for compounds which produced different effects across cell lines but also produced similar responses to other compounds on the plate, resulting in plate 16 (fig. S11A).

### Cell Lines and Culture Conditions (melanoma perturbation hit validation)

HeLa, A375, and SK-MEL-2 cells were cultured in DMEM supplemented with 10 % FCS, 1 mM pyruvate, 100 U/ml penicillin and 100 mg/ml streptomycin at 37 °C and 5 % CO_2_. All cell lines were detached for passaging with 0.05 % Trypsin at 37 °C for 15 minutes after one PBS wash and then rinsed off with DMEM. Cells were plated one day before perturbation with the indicated compounds for the indicated duration. Compounds were purchased from the vendors in Table S11.

### CellTiterGlo (CTG) assay

Cells were plated into microtiter plates (ThermoFisher Scientific #167542) as described above. 50µL CellTiterGlo substrate mix (Promega #G7573) were added to 150µL culture medium and incubated at room temperature in the dark for 10 minutes. 15µL were then transferred to white microtiter plates (Greiner Bio-One #784075) and luminescence was measured with a tecan SPARK 20M microplate reader.

### Live Cell Microscopy and Annexin-V staining

For live cell microscopy, cells were plated in fluorescence microscopy plates (ibidi #89606) and Annexin-V Alexa Fluor 647 (BioLegend #640912) was added together with the tested compounds at a 1:500 final dilution. Cells were imaged on a Leica DMi8 microscope equipped with an incubator at 37 °C and 5 % CO_2_.

### Cell Lines and Culture Conditions

Cell lines were cultured in Dulbecco’s Modified Eagle’s Medium (DMEM, Thermo Fisher Scientific, 41966-029) supplemented with 10% fetal bovine serum (FBS, Thermo Fisher Scientific), 1% Penicillin-Streptomycin (Thermo Fisher Scientific), and 1% non-essential amino acids (MEM NEAA, Thermo Fisher Scientific). The cells were maintained at 37°C in a humidified atmosphere with 5% CO2.

The following cancer cell lines were used for the screening: human lung carcinoma cell line A549, human hepatocyte carcinoma cell line HepG2, human colon carcinoma cell line HCT116, human fibrosarcoma cell line HT-1080, human colon adenocarcinoma cell line HT29, human

T-cell leukemia cell line Jurkat, human breast cancer cell line MCF7, and human osteosarcoma cell line U2OS.

The following cell lines were provided as gifts: HCT116 from Götz Hartleben, HT29 from Joel Schick, Jurkat from Daniel Krappmann, and MCF7 from Martin Selmansberger. A549, HCT116, HepG2, HT-1080, and U2OS cells were purchased from ATCC.

### Compound Screening

For the screening, 500 cells per well were seeded in 80 µl of medium in 384-well microplates (CulturPlate, PerkinElmer). After 24 hours, the cells were treated with compounds (0.4 µl per well, 5 µM final concentration) or DMSO alone as a control. The HTS platform system, consisting of a Sciclone G3 Liquid Handler (PerkinElmer), was used for plate and liquid handling. The screening plate used in this study is part of a small-molecule library with known modes of action (dissolved in DMSO, 1 mM stock solution). After 72 hours of incubation, cell viability was measured using 20 µl of CellTiter-Glo 2.0 Reagent (Promega). Luminescence signals from the CellTiter-Glo assay were detected using the EnVision 2104 Multilabel Plate Reader (PerkinElmer). The luminescence signals from the DMSO control wells were used as the baseline, representing 100% activity.

### Constructing the genetic KO experimental space for zebrafish development

Because the original experiment contained combinatorial perturbations, it was possible for us to also make combinatorial predictions during inference. However, the computational cost of the pairwise combination of all genes would have been immense, and therefore we made predictions for 1) the original 23 genetic KOs from Saunders et al, in all pairwise combination with 200 randomly selected transcription factors (Fig. 6c) and 2) the single genetic KOs of 2.1k transcription factors, as provided to us by Madeleine Duran from the Trapnell lab (Fig. 6d).

## Supplementary Notes

### Prompt syntax for Prophet

Users query Prophet with a description of the experiment in terms of categorical variables. For instance, {cell_line: A375, drug1: vemurafenib, drug2: cobimetinib, phenotype: viability} is a description of a single experiment. Each categorical label is associated with a unique vector, which is embedded into a token for the model. The model then predicts the scalar outcome associated with that experiment (e.g. 0.7).

For example, the model will take a prior knowledge embedding representing A375, prior knowledge embeddings representing vemurafenib and cobimetinib, and a learnable embedding representing viability, and will combine them to finally predict the result of the experiment, producing a scalar value that reflects the expected viability under the specified conditions.

Prophet can make a prediction for whichever phenotype it has been trained in; since the training set contains viability, gene expression, and morphological features, then Prophet is capable of making predictions for viability, gene expression, and morphological features, depending on the query. If the user further fine-tunes Prophet on proliferation data, Prophet can output predictions also about cellular proliferation.

### Clarification of usage of “phenotype”

We distinguish between the *phenotype* that is intended to be measured (e.g. gene expression, viability, cell morphology, etc.) and the *result* measured (i.e. the scalar value used to quantify the desired phenotype). For instance, the desired *phenotype* can be *cell eccentricity* and the *result* might be 0.5, assuming that 0.5 makes any sense in the selected scale to measure *cell eccentricity*. This would be model with Prophet in the following manner *Prophet*(*perturbation*, *cell state*, *cell eccentricity*) = 0. 5 being *perturbation* and *cell state* the perturbation applied to the cell state in which cell eccentricity wants to be measured.

Prophet is capable of making predictions for any modality for which it has been trained or fine-tuned on relevant data. For example, given a Cell Painting drug screening dataset such as JUMP, Prophet can predict morphology features for any compound or cell line. As described previously, Prophet generates predictions for a single phenotype at a time, as specified by the user. If the user queries cell eccentricity, *Prophet(perturbation, cell state, cell eccentricity) = 0.5*. If the user subsequently requests the IC50 value under the same perturbation and cell state, *Prophet(perturbation, cell state, IC50) = 0.7*.

### Clarification of Prophet’s input and output data

Prophet is generalizable and can, in principle, make predictions for an open-ended number of perturbation conditions. However, there are practical limitations tied to its current implementation and training data. For CRISPR perturbations, the pretrained Prophet model supports gene targets that are defined by the HUGO Gene Nomenclature Committee (HGNC) for human and mouse genomes. For unseen cell lines, predictions are limited to those for which transcriptomic profiles are available in the CCLE dataset, since cell state embeddings are derived from these profiles. Users may extend Prophet to other perturbations or cell states by retraining the model with appropriate input embeddings.

Prophet is a univariate regression model, so it operates on scalar targets. For high-dimensional data such as transcriptomics or Cell Painting features, which are typically represented as vectors, we decompose the task into multiple scalar regression problems—one per feature. However, all scalar predictions are made using the same function: Prophet(cell, perturbation, feature) → feature value. This means that the model shares parameters across all features and conditions, enabling it to learn shared representations and exploit statistical dependencies between features during training.

## Supplementary Figures

**Fig. S1.**
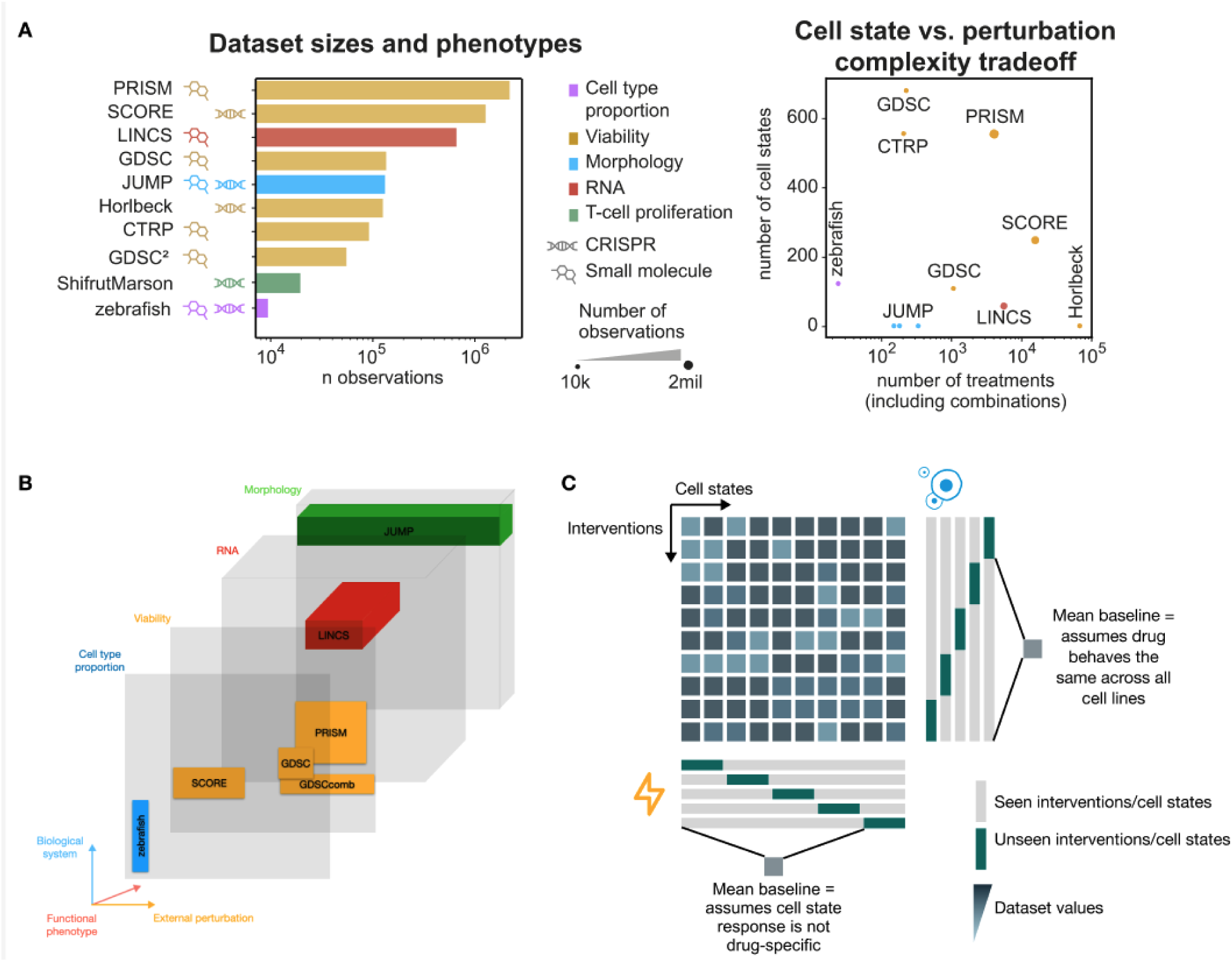
Overview of the datasets and additional performance numbers. A) The datasets vary both in number of cell states and number of perturbations, and include drug and genetic perturbation. Not indicated: GDSC^2^, Horlbeck, and zebrafish are combinatorial. B) Approximate representation of the coverage of the dataset compendium in experimental space. We gathered 10 perturbational datasets that encompass up to 4 different phenotypes: IC_50_/viability, RNA measurements, cell type proportion and cell morphology, with varying amounts of overlap. In total, there are more than 4.7 million experiments. C) Schematic of how 5-fold cross-validation is constructed for a single seed, resulting in ten runs. This is repeated for 3 seeds. The grey and blue tiles each represent one value in the experimental matrix of a dataset. The mean baseline estimates that the effect of an perturbation in an unseen cell state is the average value of that perturbation in all previously seen states. Conversely, it estimates the effect of an unseen perturbation on a cell state is the average of all seen perturbations in that same cell state. Naturally, it changes depending on the test set used.

**Fig. S2.**
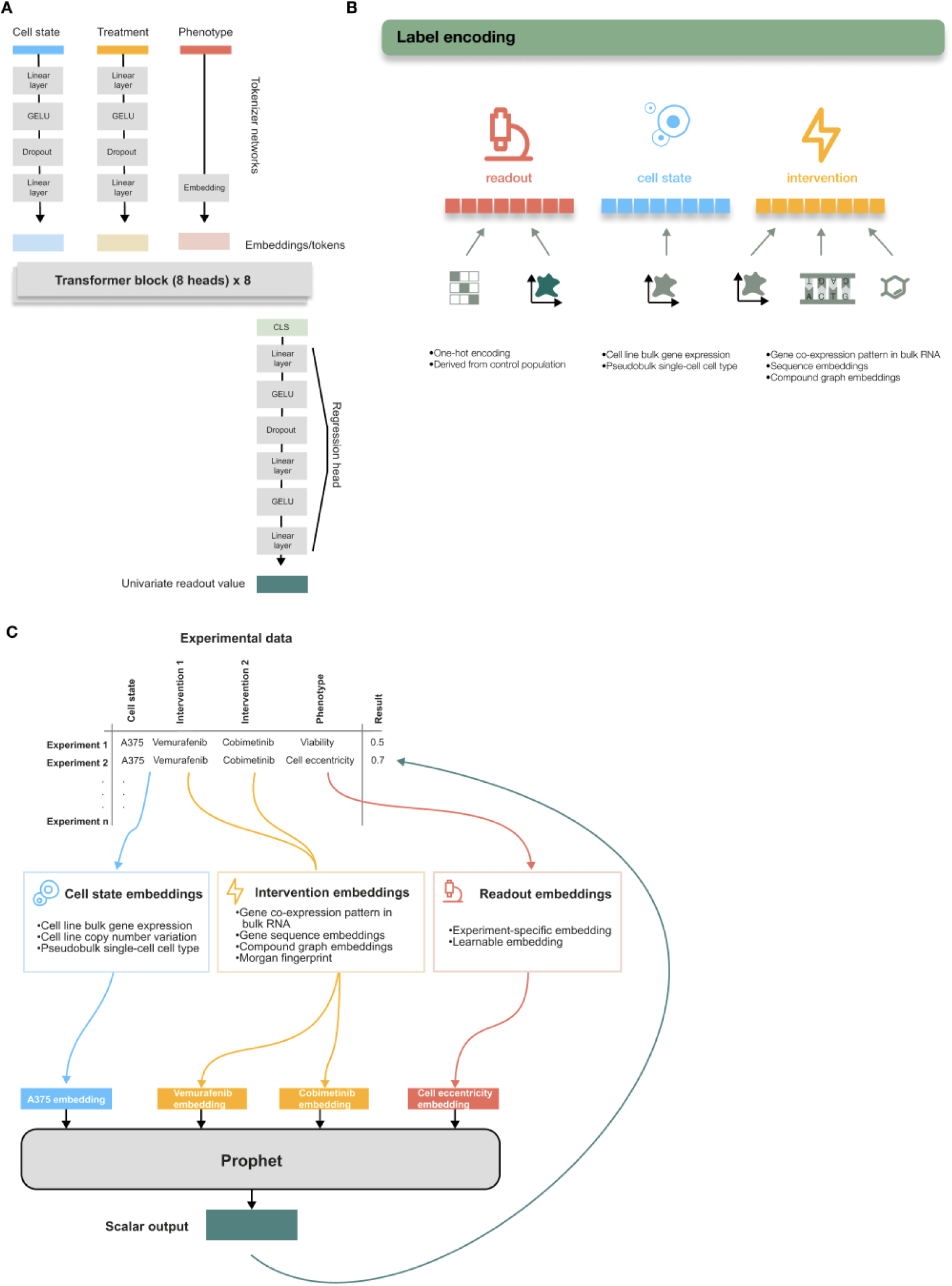
Prophet’s inputs, outputs, and architecture. A) Prophet is a transformer-based model consisting of MLPs that serve as tokenizers for categorical input variables, 8 transformer blocks with 8 attention heads each, and a MLP as a regression head that outputs the predicted phenotypic value. B) Cell state, perturbation, and phenotype are encoded as vector representations when input into the model. C) Schematic of how each experiment is represented in a dataframe format. Embeddings are built of each element of the experimental set and then feeded into Prophet, which predicts the resulting scalar value.

**Fig. S3.**
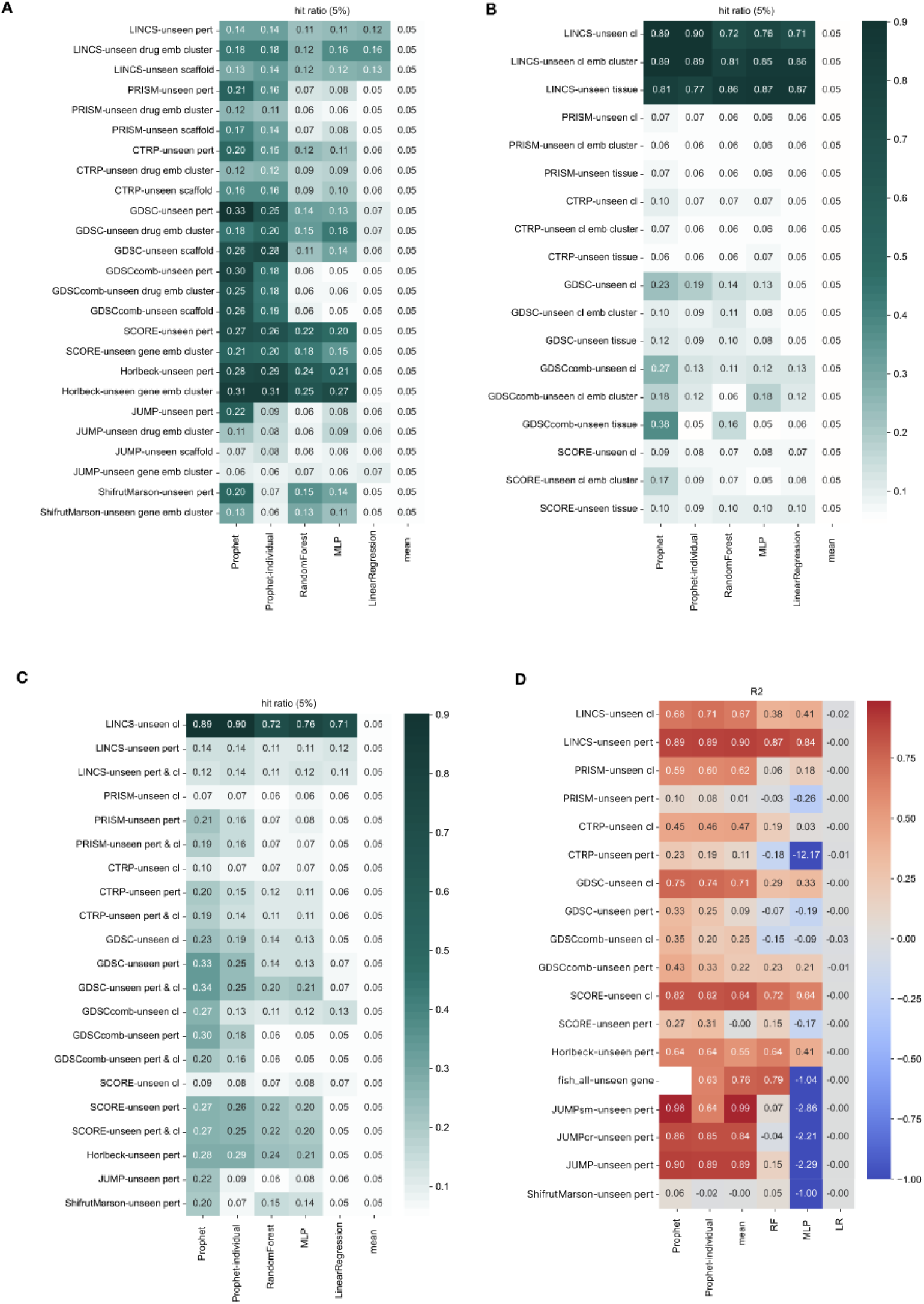
Performance tables using harder partitions of each dataset. A) Extrema 5% hit ratio for all datasets and three splits (where applicable): unseen cell lines (’leave_cl_out’), unseen perturbations (’leave_genes_out’), and both unseen cell line and perturbation (’leave_both_out’). Excluding the LINCS dataset, Prophet performs better in all but one case. B) Extrema 5% hit ratio for variations of the unseen cell line partition. Prophet performs better in all but one case. C) Extrema 5% hit ratio for variations of the unseen perturbation partition. Prophet performs better in 20/25 cases. D) The same table as in fig. S5D but reporting R2 score.

**Fig. S4.**
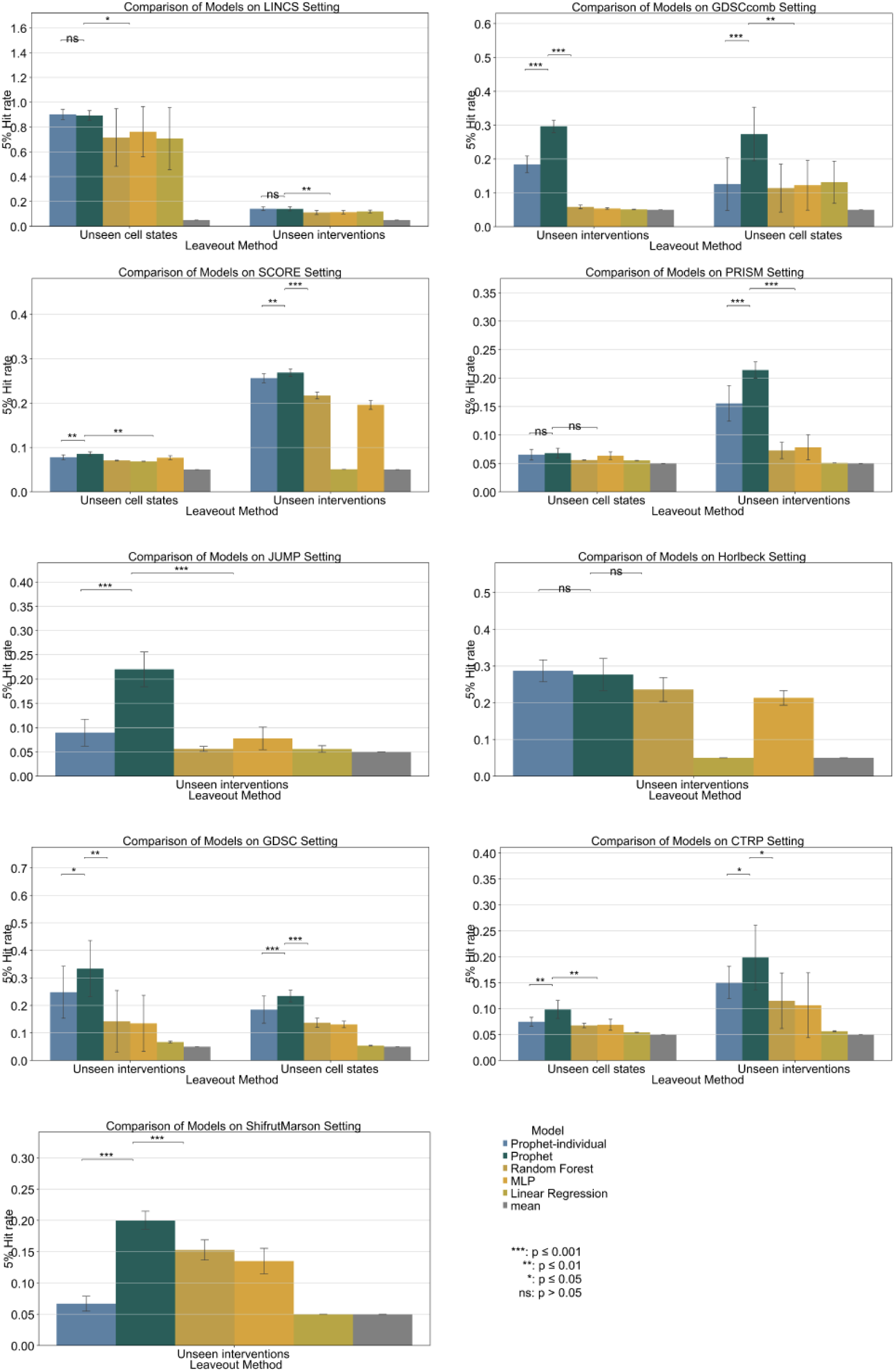
Comparative performance of Prophet and baseline models across benchmark settings. Top 5% hit ratio for Prophet, Prophet-individual, and baseline models. Each subplot corresponds to a specific dataset and displays performance under two challenging generalization regimes: unseen cell states and unseen perturbations. Prophet consistently outperforms baseline models across most settings, particularly under the harder unseen perturbations condition. Statistical significance is denoted using Wilcoxon signed-rank test comparisons against Prophet, with thresholds: ***p ≤ 0.001, **p ≤ 0.01, *p ≤ 0.05, ns: not significant. Results are aggregated over three random seeds and five cross-validation splits, and error bars represent standard error of the mean.

**Fig. S5.**
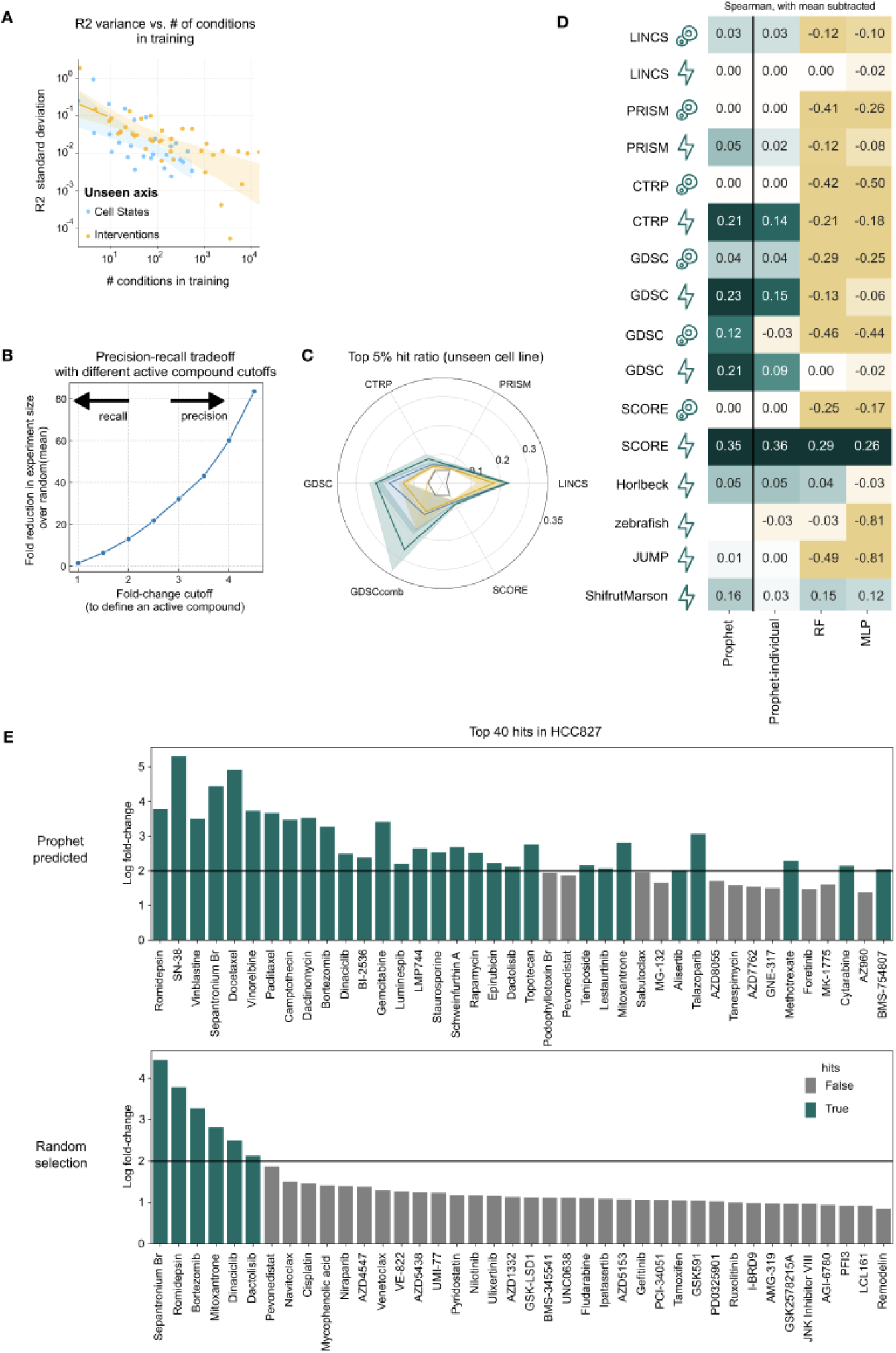
Mean baseline and evaluation schema. A) The variance in the R2 score obtained by Prophet is reduced as more parts of the experimental space - more cell states and more perturbations - are sampled during training. B) Precision–recall tradeoff as a function of fold-change threshold used to define logFC hits. Higher thresholds represent precision in finding hits with fewer experiments, lower thresholds represent ability to recall hits using the top experiments. C) Same radar plot as in Fig. 2E but for unseen cell lines - top 5% hit ratio across held-out cell lines for Prophet evaluated on datasets with more than ten cell lines. D) The same performance table as in fig. S5D, but reporting Spearman correlation instead of hit ratio. E) Example of how using Prophet to select compounds affecting viability in HCC827 cells results in discovery of more hits over random selection. Top panel is the top 40 compounds as selected by Prophet, bottom panel is a random selection of 40 compounds. X-axis represents the real log-fold change in viability over DMSO perturbation in the same cell line.

**Fig. S6.**
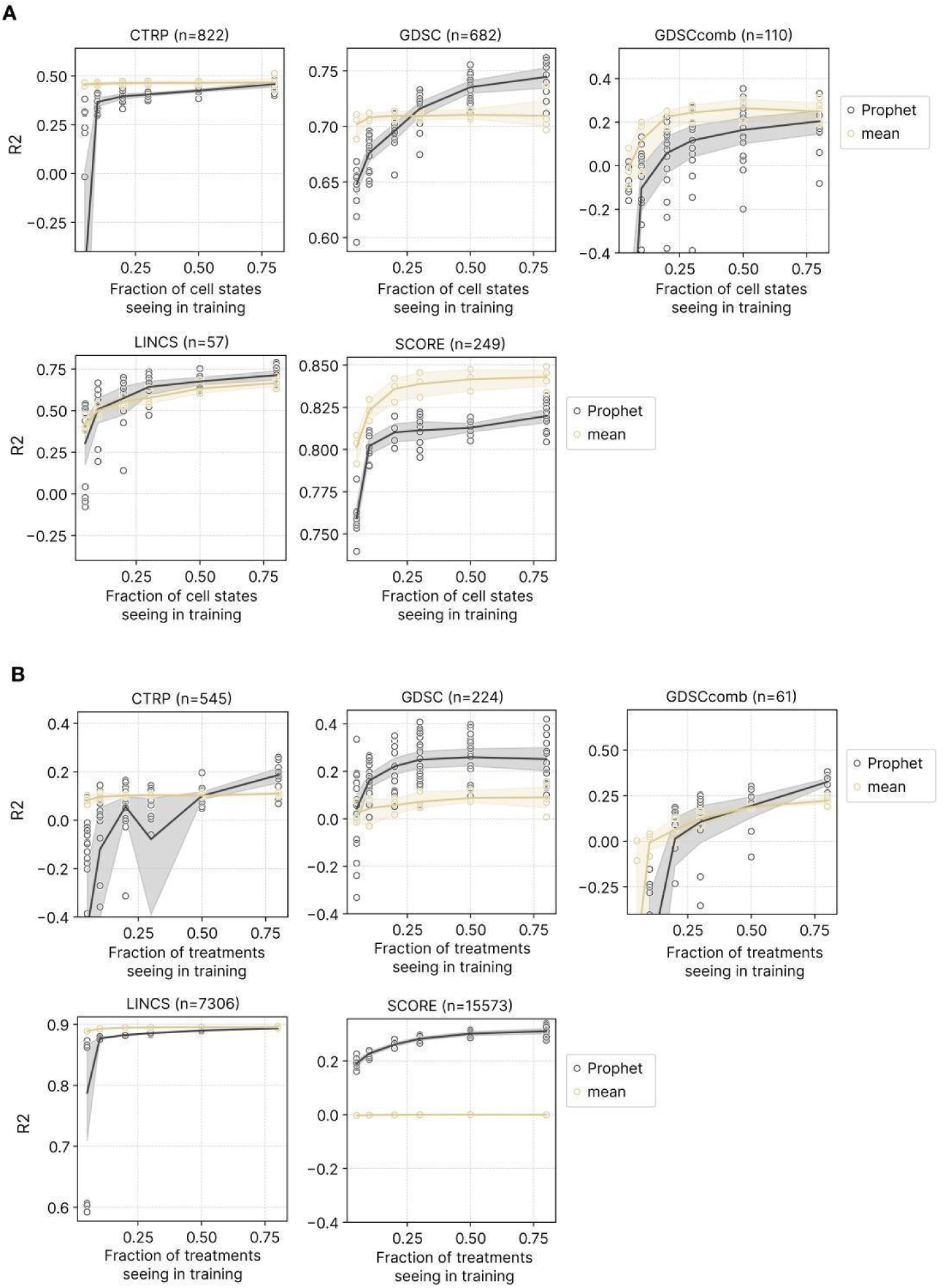
Prophet-individual models’ performance as a function of the number of conditions seen during training. The more the experimental space is mapped during training, the better Prophet’s performance. However, the concrete pattern is dataset dependent as different datasets capture different amounts of variation. A) Performance of Prophet-individual models along the cell line axis. B) Performance of Prophet-individual models along the perturbation axis.

**Fig. S7.**
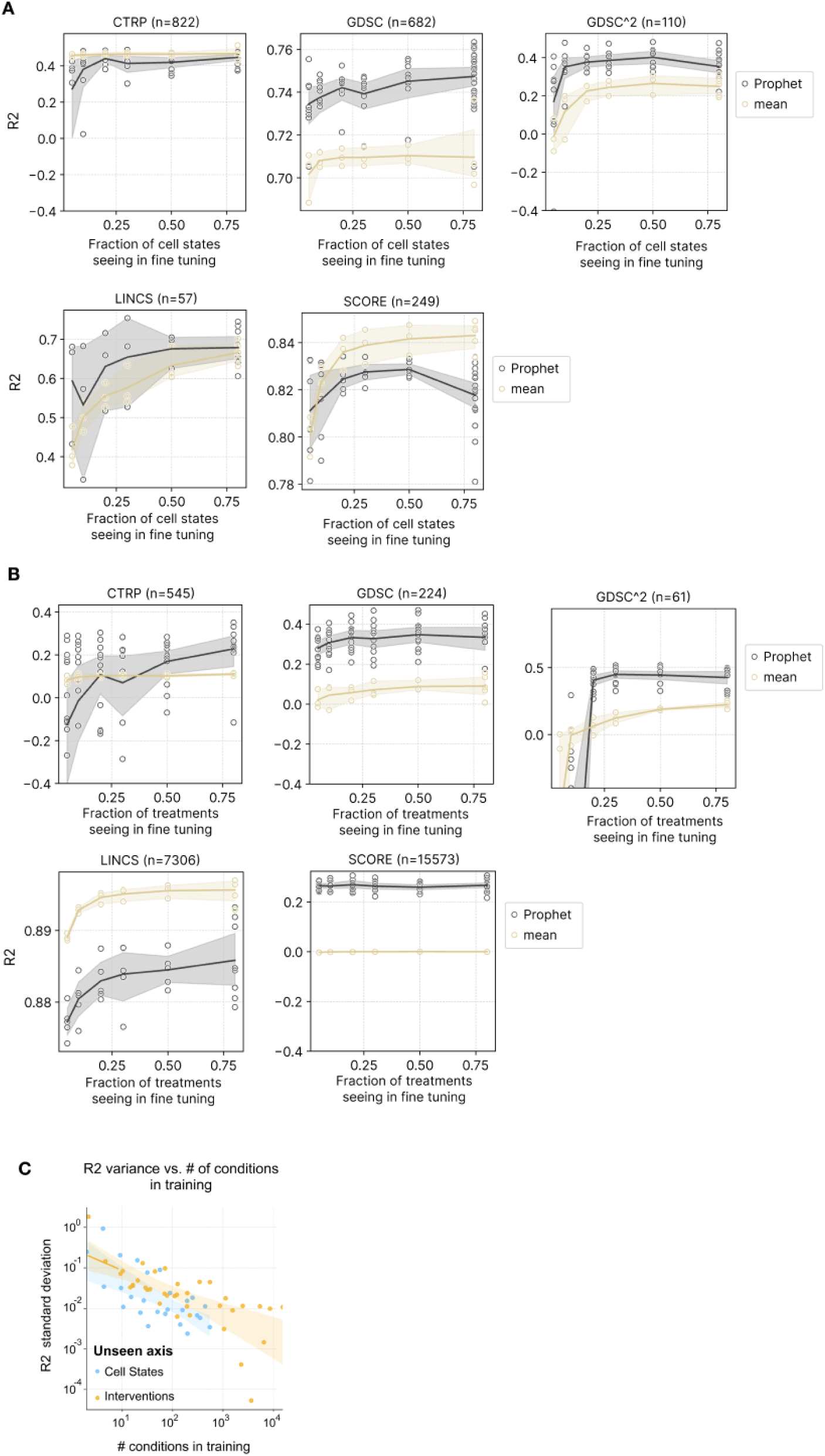
Prophet models’ performance as a function of the number of conditions seen during fine tuning. It can be seen that performance is more robust when fine tuning. As a remark, all models here are fine tuned for just 20,000 steps, independently of the size of the dataset. A) Performance of Prophet along the cell line axis. B) Performance of Prophet along the perturbation axis. C)

**Fig. S8.**
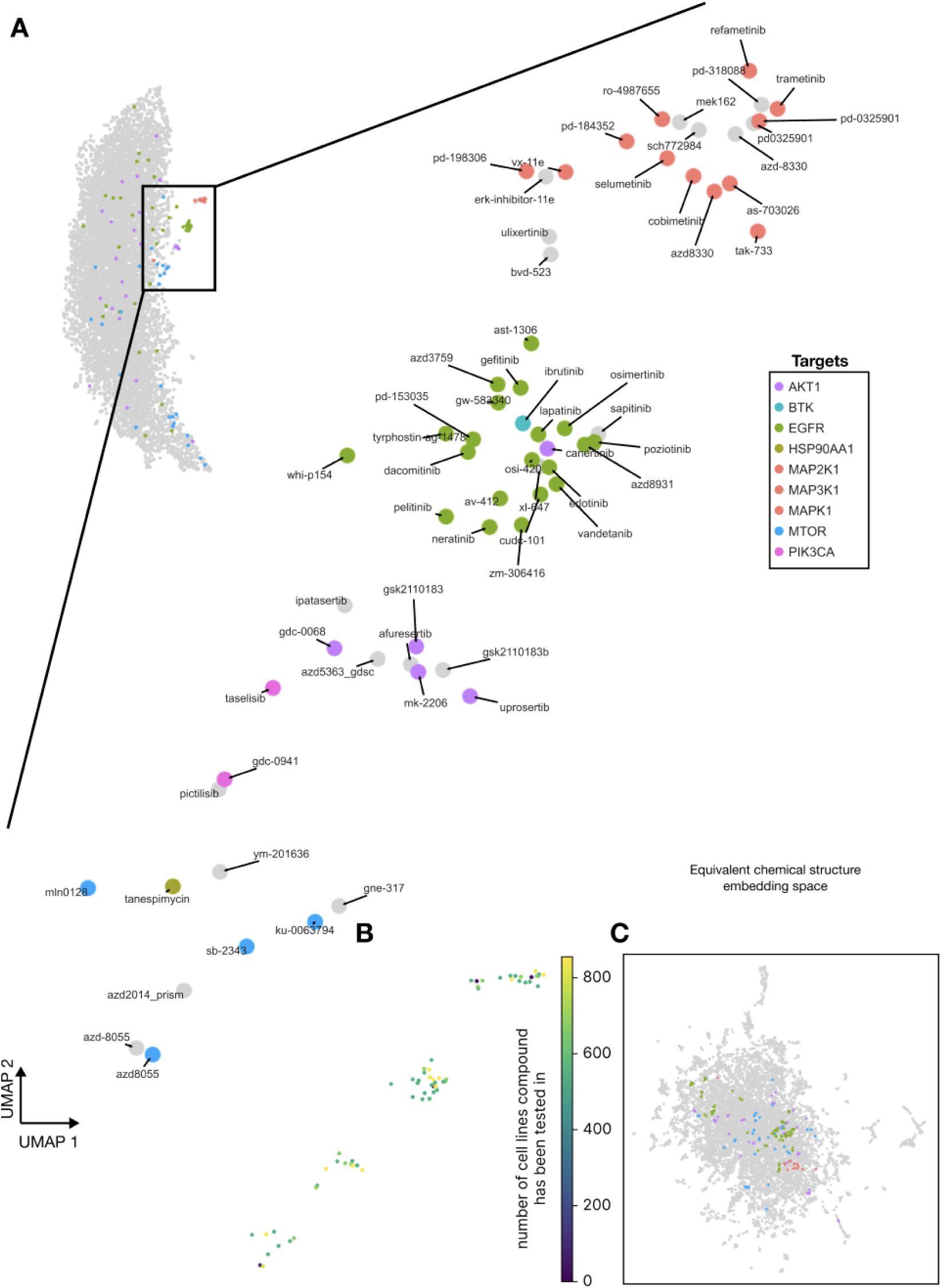
Detail of the clusters found in the viability Prophet experimental space. A) Drugs are annotated by their automatically fetched annotations from pertpy. Compounds in grey did not have a putative target annotation in the database, but may still be annotated by existing literature. B) The number of unique cell lines in which a compound has been tested, for the 4 zoomed-in clusters. The fewest is 0 and the most is 829. C) Chemical structure embedding for the same set of compounds as shown in A, with the same 4 clusters colored.

**Fig. S9.**
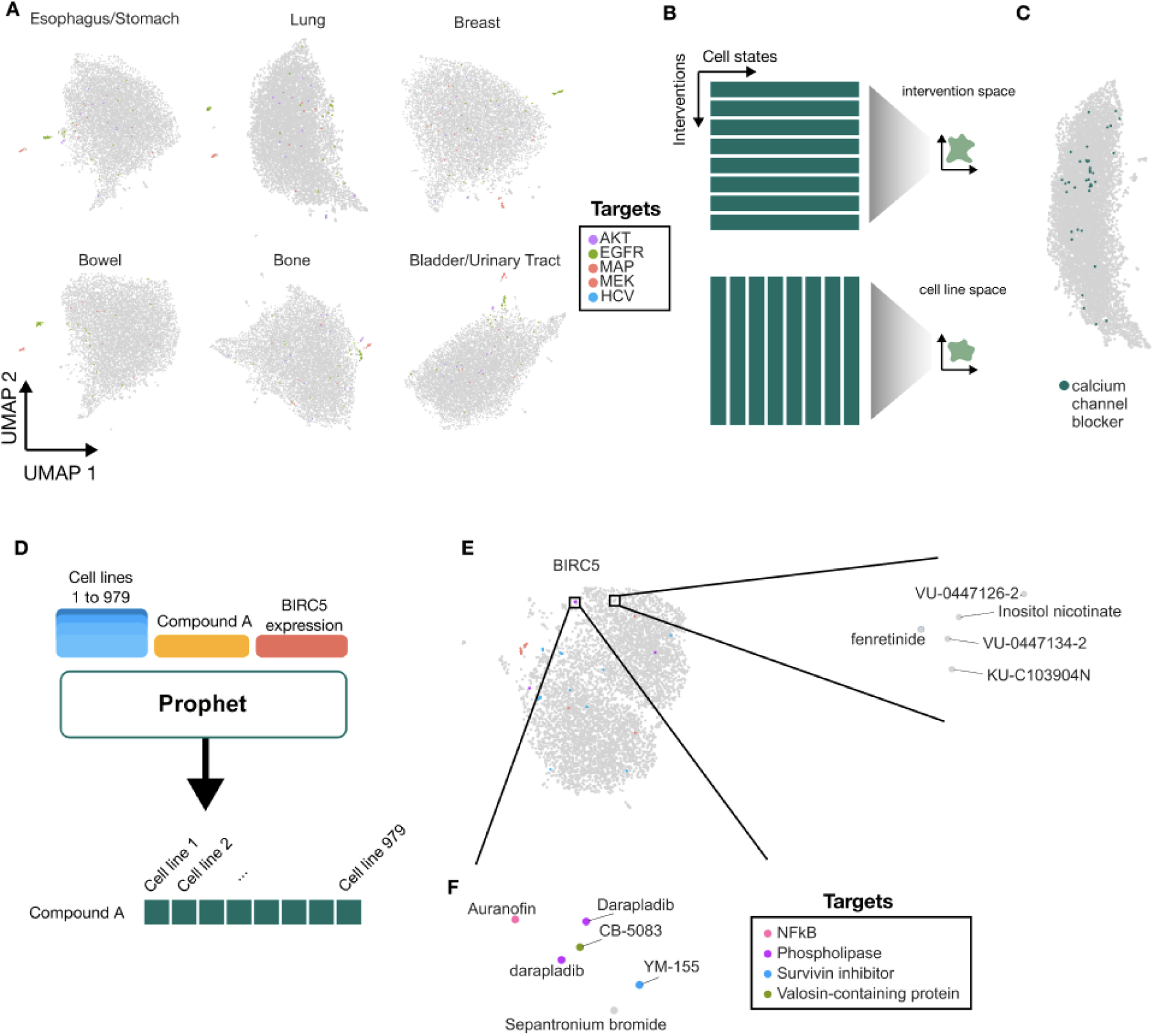
Experimental space construction and additional examples. A) The 8620 compounds depicted in Fig. 3B, but subset to cancer cell lines of different tissue origins. Different tissues demonstrate different response patterns to compounds. B) Graphical depiction of how cell line spaces and perturbation spaces are generated. Each teal rectangle represents a vector which is embedded as a 2D point using UMAP. C) Calcium channel blockers in the experimental space shown in Fig. 3B. Calcium channel blockers do not have a cohesive activity fingerprint across all tissue types, in contrast to when subset to brain and CNS-derived cancers. D) Schematic of how the BIRC5 experimental space is constructed. E) The experimental space of BIRC5 expressed across 979 cell lines after perturbation with 8620 compounds. This is the BIRC5 RNA expression space equivalent of Fig. 3B, which depicts IC_50_ space instead. A potentially interesting cluster of unknown compounds is annotated (right). F) A cluster of compounds with some putative regulators of BIRC5 (survivin).

**Fig. S10.**
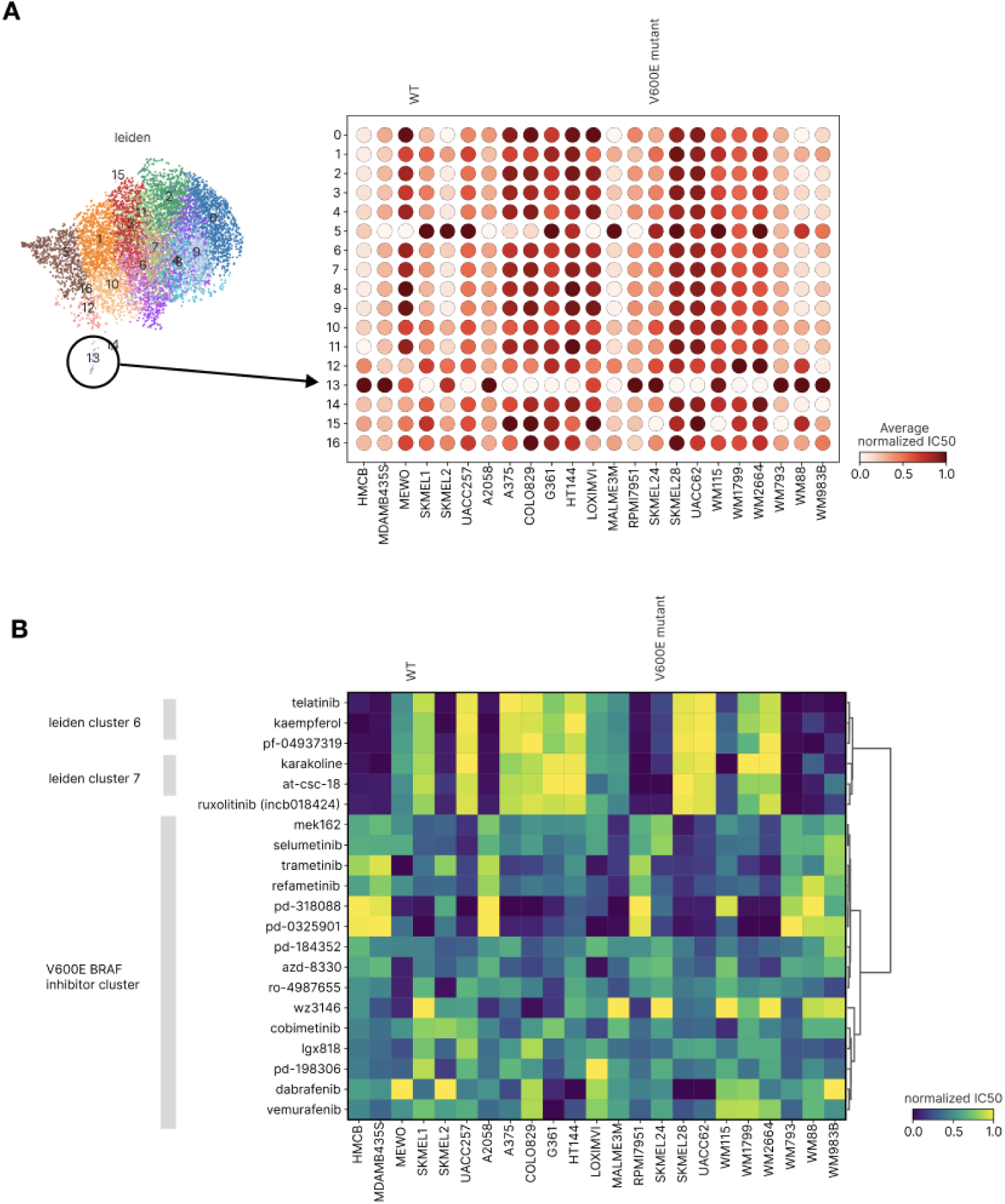
In silico modeling of perturbation effect in V600E *BRAF* mutant and wild-type cell lines. A) The space of 23 V600E mutant or wild-type melanoma cell lines, as in Fig. 3D, colored by leiden clustering. The dotplot depicts the average IC_50_ value across compounds per cluster, with darker clusters therefore indicating more resistance to the compounds in that cluster. The left 6 cell lines are putatively wild-type and remaining cell lines are putatively V600E *BRAF* mutants. B) Examples of compounds from A. Prophet predictions for all compounds from leiden cluster 13 as well as 3 random compounds selected from leiden clusters 6 and 7 for contrast. Prophet predicts a distinctly different pattern of behavior for compounds in cluster 13, which also contain known V600E *BRAF* inhibitors.

**Fig. S11.**
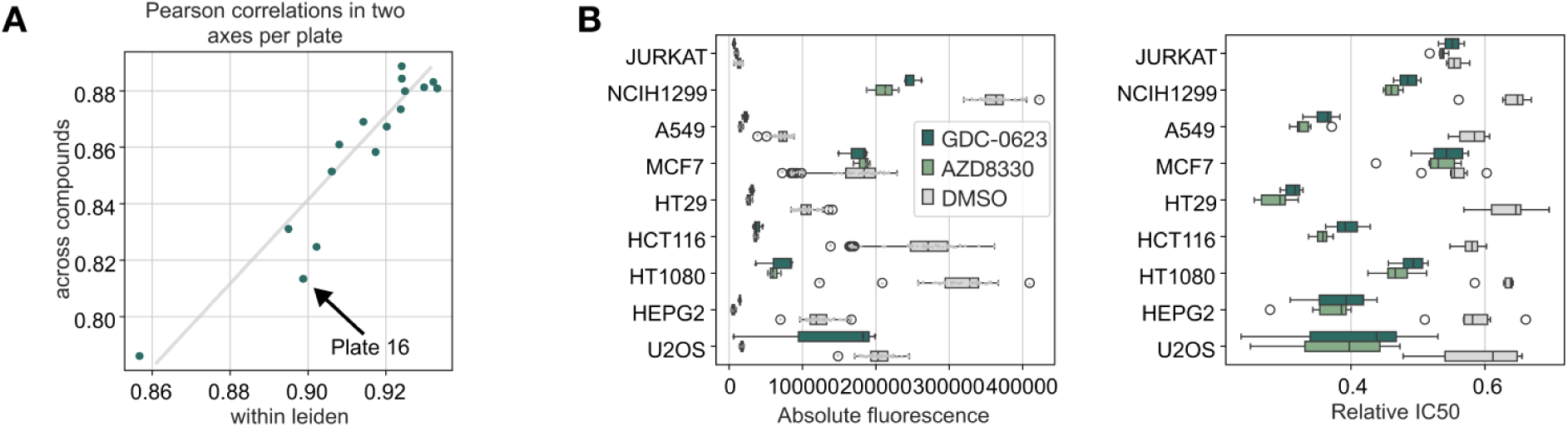
Experimental design and in vitro validation. A) Metrics used to select one out of 16 possible plates in an in vitro drug screening setup. First, Prophet was used to predict the outcome of each 352-compound plate in 9 cell lines. X-axis reports the average pairwise Pearson correlation between compounds within the same leiden cluster. Y-axis reports the average pairwise correlation between all compounds. We optimized for low correlation across compounds and high correlation within cluster (distance from the regression line). B) The same plots as for Fig. 5E, but including all cell lines. Prophet also correctly predicts that AZD8330 has, for most cell lines, a stronger effect on viability than GDC-0623.

**Fig. S12.**
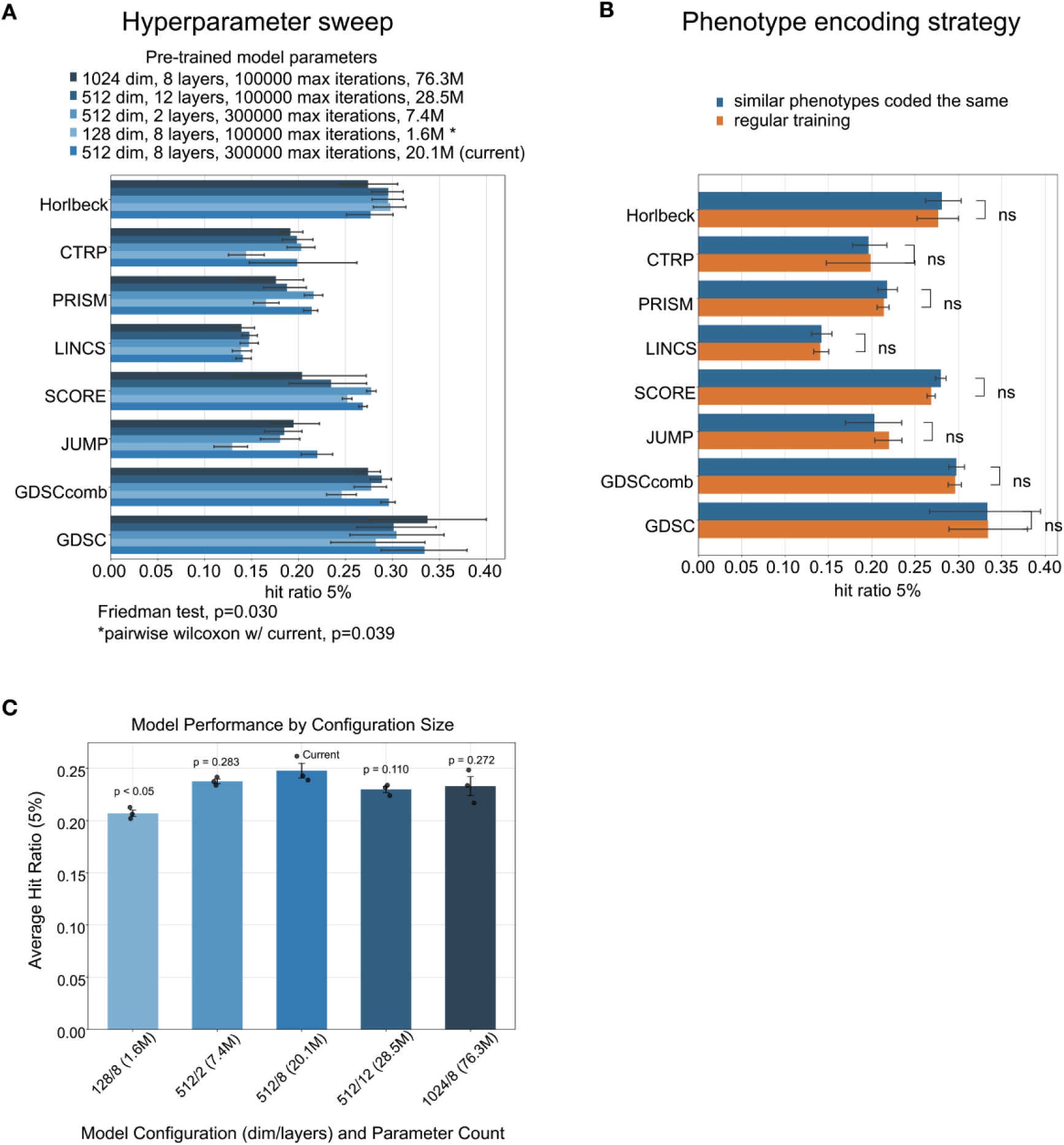
Evaluation of Prophet’s robustness to pretraining configurations and phenotype modeling strategies. A) Performance of various pretraining configurations on 5% hit ratio across datasets, showing Prophet’s stability across different pre-training architectural choices (e.g., latent dimension, depth, iteration count) except in the case of too few parameters(red). B) Effect of how the phenotype label is represented on generalization to unseen perturbations. Coding similar phenotypes the same does not change overall performance compared to regular training. C) 5% hit ratio across Prophet pretraining of increasing size. The current Prophet pretraining configuration (512/8, 20.1M parameters) happens to achieve the highest average hit ratio, though differences with other large models are not statistically significant. A small model (e.g., 128/8) shows significantly lower performance (*p* < 0.05). Error bars represent standard error of the mean across seeds.

## Supplementary Tables

**Table S1.** Dataset overview. Shown are the metrics of the datasets employed. The perturbations are divided into CRISPR or drugs and combinations or singletons. The phenotypes range from IC_50_ of viability to cell type proportions. In the case of JUMP and LINCS, the number of phenotypes is larger than 1 since there are several cell painting features predicted and a total of 50 selected genes. The number of combinations represents the number of unique combinations of perturbations. It is different from the number of perturbations just in the case of datasets with combinations of perturbations. The number of experiments represents the total number of datapoints in the dataset. Notice that it is not the number of combinations times the number of cell states since some datasets do not measure the entire experimental matrix. The sparsity column denotes how completely the experimental matrix was filled in for seen conditions.

| Name | perturbation | Readout type | # of readouts | # of cell states | # of perturbations | # of combinations | # of experiments | Sparsity | Included in Prophet |
| --- | --- | --- | --- | --- | --- | --- | --- | --- | --- |
| **Horlbeck et al.**<sup>242424</sup> | CRISPR combination | viability | 1 | 2 | 367 | 67384 | 126037 | 0.935 | Yes |
| **GDSC**<sup>212121</sup> | Drug singleton | ln(IC50) | 1 | 682 | 224 | 224 | 135106 | 0.884 | Yes |
| **GDSC**<sup>222222</sup><sub>2</sub> | Drug combination | ln(IC50) | 1 | 110 | 61 | 1195 | 61795 | 0.470 | Yes |
| **cpg-0000 jump**<sup>888</sup> | Drug/CRISPR singleton | Cell painting features | 200 | 2 | 404 | 404 | 161600 | 1.000 | No |
| **LINCS**<sup>202020</sup><sub>0</sub> | Drug singleton | RNA (fluorescence) | 42 | 57 | 7306 | 7306 | 1491002 | 0.085 | Yes |
| **PRISM**<sup>111</sup> | Drug singleton | Log2FC | 1 | 556 | 2065 | 2065 | 1121973 | 0.977 | Yes |
| **SCORE**<sup>333</sup> | CRISPR singleton | essentiality (CERES) | 1 | 249 | 15573 | 15573 | 1279146 | 0.330 | Yes |
| **Saunders et al.**<sup>252525</sup> | CRISPR singleton/combination | celltype proportions | 1 | 620 | 24 | 24 | 9370 | 0.630 | No |
| **CTRP**<sup>232323</sup> | drug singleton | IC50 | 1 | 822 | 545 | 545 | 357865 | 0.799 | Yes |

**Table S2.** Overview of architecture hyperparameters used for the Prophet-individual models. Shown are the model hyperparameters used for Prophet-individual models. The dimensionality of the tokenized embedding, number of transformer blocks and the learning rate were the most influential hyperparameters, while other factors had minimal impact.

| Hyperparameter | Value |
| --- | --- |
| Warmup steps | 10000 |
| Batch size | [2048, 1024, 512, 256] |
| Precision | float32 |
| Embedding dimensionality | [512, 256, 128] |
| Transformer blocks | [4,2] |
| Attention heads | [1,2,4] |
| Maximum learning rate | [.0001, .0005] |

**Table S3.**
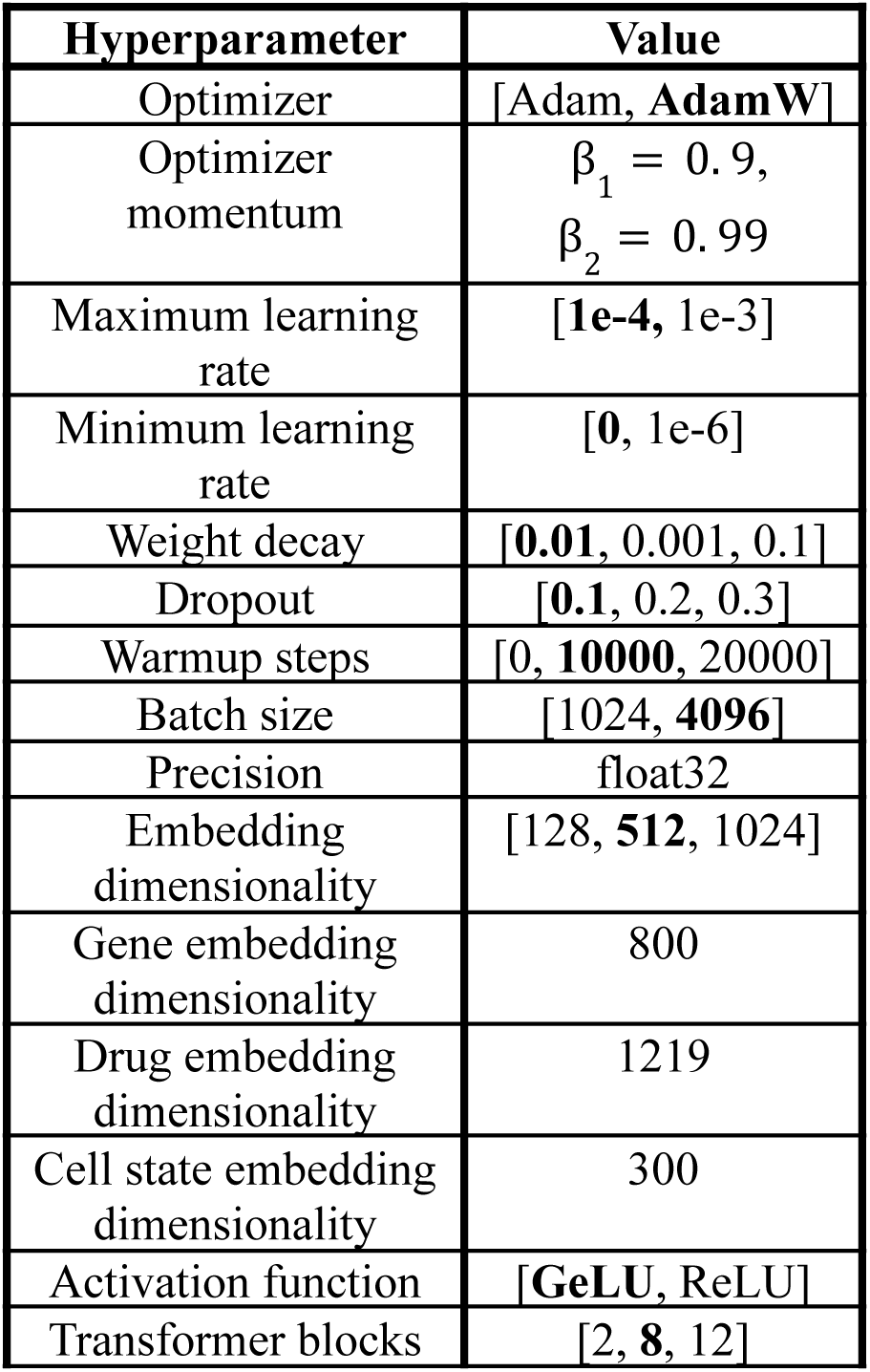
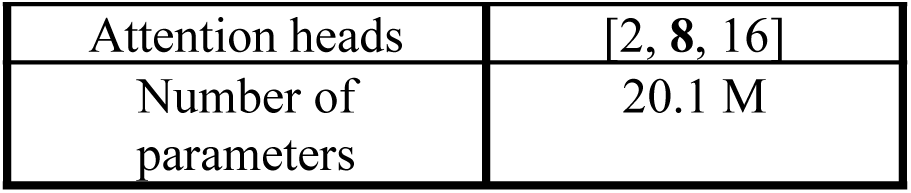
Overview of pretraining architecture hyperparameters used for the final Prophet pretraining model. Shown are the model hyperparameters used for Prophet’s pretraining, in bold are the hyperparameters employed for the final pretrained models. In general, the activation function, the dropout, minimum learning rate and batch size choices did not have a major effect on the performance. Five-fold cross validation across three seeds and two data splits (unseen cell states and unseen perturbations) required generating 30 pretrained models for a total of approximately 400 GPU hours (A100). Instead, for most hyperparameters we run smaller models. We ran, however, large Prophet models hyperparameter searches with alternative transformer configurations and different number of parameters that showed the role the embedding dimensionality of the model as a main driver of performance and that scaling beyond ∼20M parameters in the pre-trained setting offered no performance benefit (fig. S12).

**Table S4.** Statistical metrics of the SCORE dataset across the cell line axis. Shown are the Pearson correlation scores obtained on the SCORE dataset evaluating the replicates along the cell line axis. The Pearson correlations are computed between random replicates, between the replicates that belong to a cell line (within a cell line) and across cell lines once the replicates are all aggregated (across cell lines). In each case, they are computed for different scenarios: no replicate is aggregated, just the perturbational replicates are aggregated, just the cell line replicates are aggregated, and all replicates are aggregated. Importantly, the Pearson correlation between random replicates once the perturbational replicates are aggregated (0.674) is bigger than the Pearson correlation between replicates of a same cell line (0.615). This means that the cell line-specific information gets lost once the replicates are aggregated and, hence, one can not distinguish a random replicate from a real one.

| **Pearson correlation** | **No replicates aggregated** | **Perturbation replicates aggregated** | **Cell line replicates no aggregated** | **Perturbation replicates no aggregated** | **Cell line replicates aggregated** | **All replicates aggregated** |
| --- | --- | --- | --- | --- | --- | --- |
| **Between Random replicates** | 0.481 | 0.674 |  | - |  | - |
| **Within cell line** | 0.615 | 0.820 |  | - |  | - |
| **Across cell lines** | - | - |  | 0.625 |  | 0.755 |

**Table S5.** Statistical metrics of the SCORE dataset across the perturbation axis. Shown are the Pearson correlation scores obtained on the SCORE dataset evaluating the replicates along the perturbation axis. The Pearson correlations are computed between random replicates, between the replicates that belong to an perturbation (within perturbation) and across perturbations once the replicates are all aggregated (across perturbations). In each case, they are computed for different scenarios: no replicate is aggregated, just the perturbational replicates are aggregated, just the cell line replicates are aggregated, and all replicates are aggregated. Importantly, the Pearson correlation between random replicates once the cell line replicates are aggregated (0.046) is lower than the Pearson correlation between replicates of a same perturbation (0.125). This means that the perturbation-specific information does not get lost once the replicates are aggregated and, hence, one can distinguish a random replicate from a real one. Therefore, the mean estimator can be outperformed since the experiment retains biological signals.

| Pearson correlation | No replicates aggregated | Perturbation replicates aggregated | Cell line replicates no aggregated | Perturbation replicates no aggregated | Cell line replicates aggregated | All replicates aggregated |
| --- | --- | --- | --- | --- | --- | --- |
| Between random replicates | 0.029 | - |  | 0.046 |  | - |
| Within perturbation | 0.125 | - |  | 0.208 |  | - |
| Across perturbations | - | - |  | 0.106 |  | 0.131 |

**Table S6.** Enrichment of known mechanisms of action in four clusters in small molecule IC_50_ space. Each of the four distinct clusters shown in Fig. 3B describe a unique cell proliferation pathway. To demonstrate that the mechanism of action enrichment is significant, we report three statistics on the annotations in each of cluster: 1) the fraction of the small molecules in the cluster which are known to have the indicated mechanism of action 2) the fraction of all small molecules with that annotation which are in the cluster 3) p-value (Fisher exact test, Supplementary Methods). Values which were smaller than 10^−7^ are reported as 0.

| Primary mechanism of action | frac of cluster | n in cluster/all | p-value |
| --- | --- | --- | --- |
| AKT inhibitor | .36 | .27 | 0 |
| MEK inhibitor | .55 | .61 | 0 |
| EGFR inhibitor | .71 | .52 | 0 |
| mTOR inhibitor | .33 | .19 | 0 |

**Table S7.**
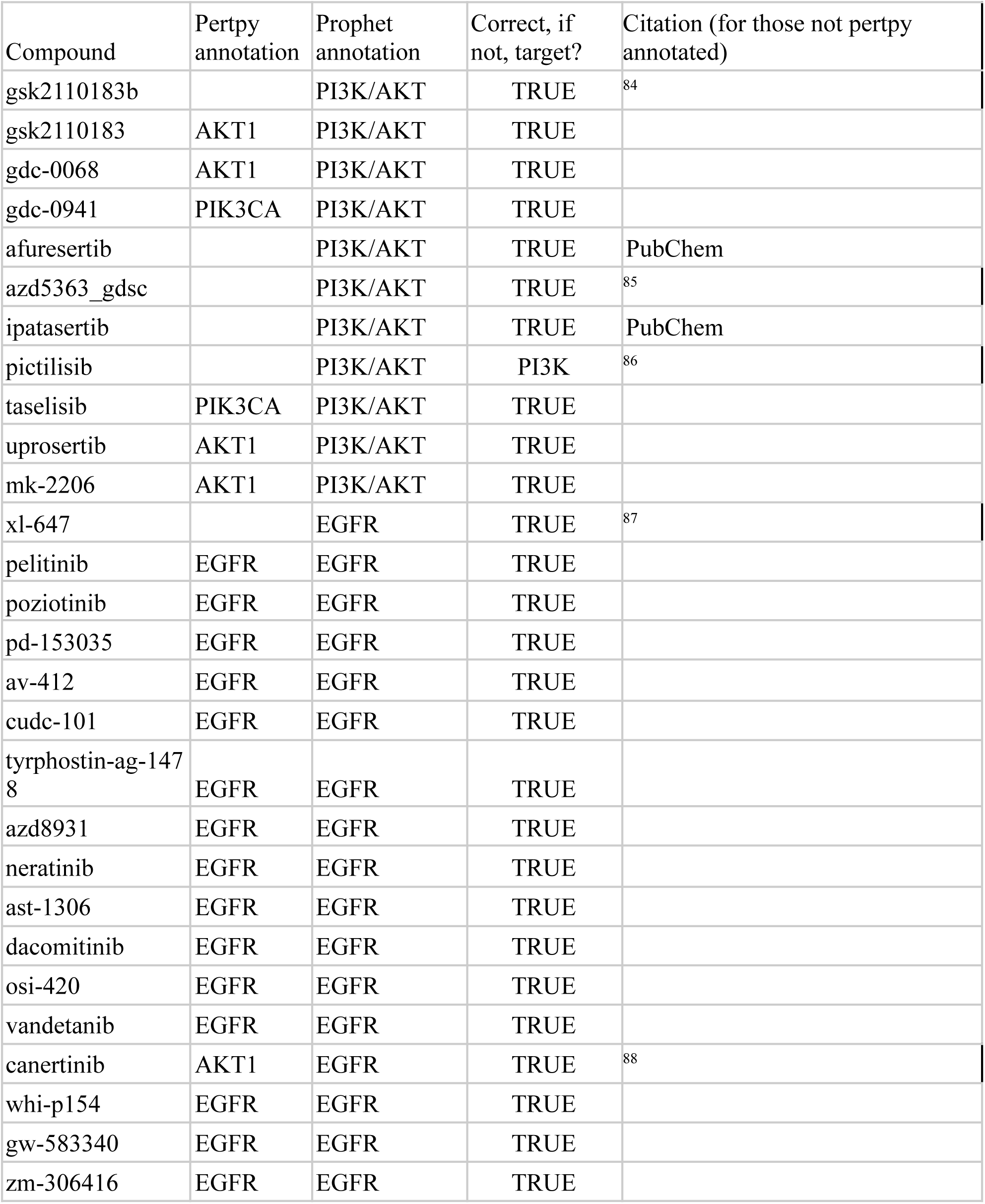
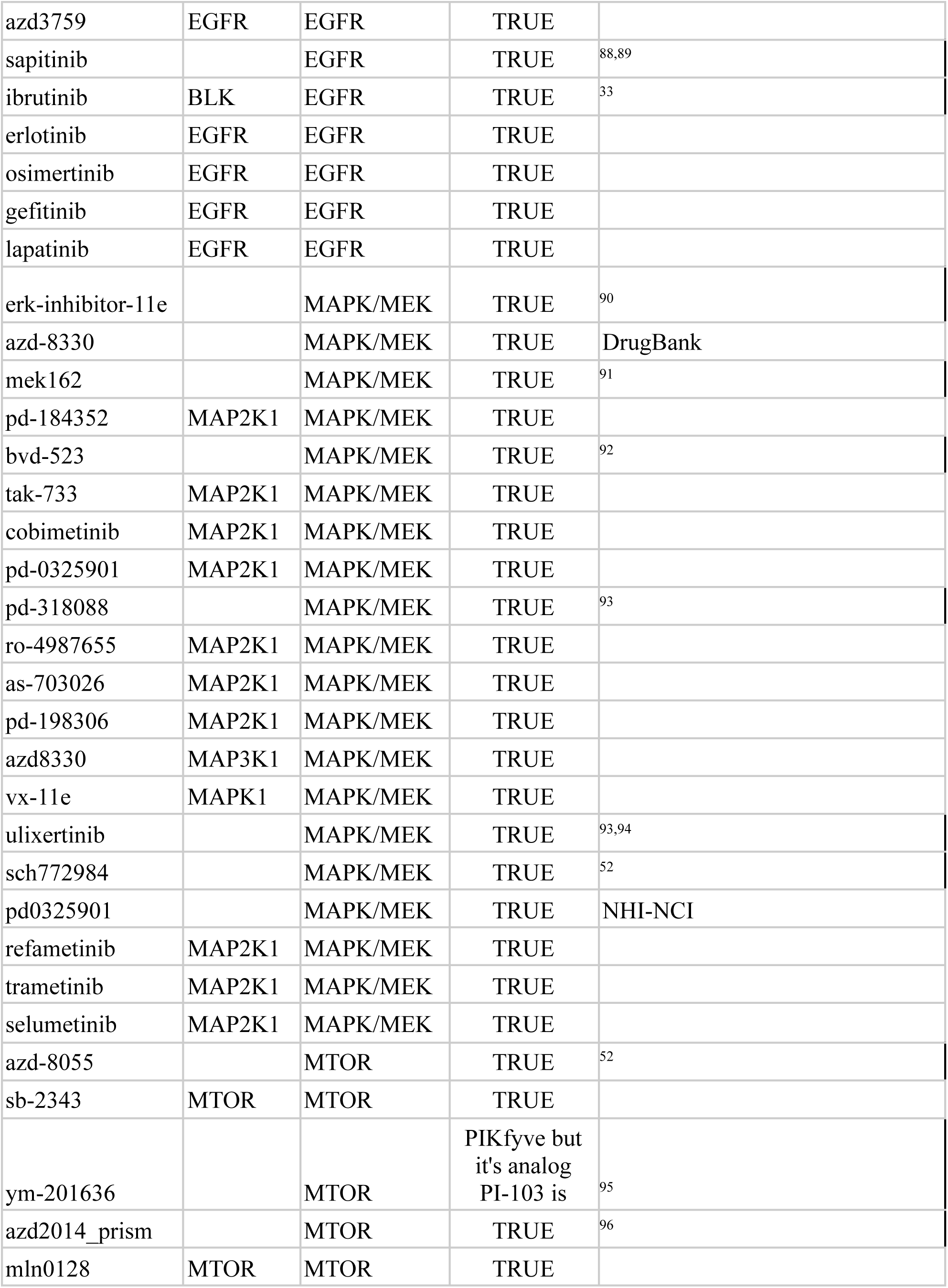
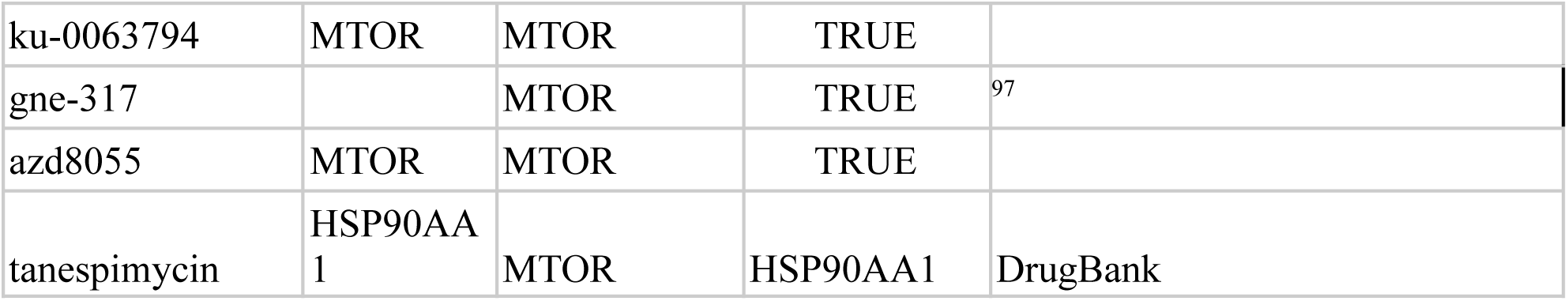
Compounds in four viability-critical pathways. Compounds which were not automatically annotated by pertpy were manually annotated. The appropriate citation is referenced in the final column.

**Table S8.** Compounds in a cluster containing V600E *BRAF* melanoma perturbations. Additional information citing each compound as indicated for either melanoma or as a cancer perturbation is referenced in the final column. The vast majority of compounds have literature backing as chemotherapies.

| Compound | Other names | Known V600E inhibitor | Clinically approved | Citation re: activity |
| --- | --- | --- | --- | --- |
| PD-0325901 | Mirdametinib | --- | Phase II, neurofibroma | <sup>98</sup> |
| PD-184352 |  | --- | No | <sup>99</sup> |
| Dabrafenib |  | TRUE | Yes |  |
| PLX-4720 |  | TRUE | No | <sup>100</sup> |
| Vemurafenib | PLX-4032 | TRUE | Yes, V600E BRAF melanoma | <sup>101</sup> |
| SCH772984 |  | TRUE, also for NRAS | No | <sup>102</sup> |
| AS-703026 | Pimasertib | TRUE | Phase II, NRAS melanoma | Only in combination with vemurafenib <sup>103</sup> |
| BVD-523 | Ulixertinib | TRUE | Phase II, GI cancer | <sup>104,105</sup> |
| Cobimetinib |  | TRUE | Yes |  |
| MEK162 | Binimetinib | TRUE | Yes, for V600E BRAF cancer |  |
| Trametinib |  | TRUE | Yes, V600E BRAF melanoma |  |
| MK-0752 |  | Known that is isn't selective | Phase I, various non-melanoma cancers | <sup>106</sup> |
| VX-11E |  | Known that it is broadly toxic | No | <sup>105</sup> |
| TAK-733 |  | Known that it isn't selective | Phase I | <sup>107</sup> |
| Selumetini<br>b |  | True, but it is<br>NOT a V600E<br>melanoma<br>inhibitor | Yes, for neurofibroma | 108 |
| WZ3146 |  | FALSE | No | No main reference |
| AZD-8330 |  | FALSE | Phase I, solid tumor | 109 |
| PD-198306 |  | FALSE | No |  |
| PD-318088 |  | FALSE | Preclinical at Pfizer | 109 |
| Refametini<br>b |  | FALSE | Phase II, hepato carcinoma | 110 |
| RO-498765<br>5 |  | FALSE | Phase I | 111 |

**Table S9.** The nine cell lines available for in vitro validation. Including known mutation statuses and tissue of origin.

| Cell Line | Gene Mutations | Tissue of Origin |
| --- | --- | --- |
| JURKAT | TP53, NOTCH1, PTEN | T-cell Leukemia (Blood) |
| NCI-H1299 | TP53 | Lung Carcinoma |
| A549 | KRAS, TP53, STK11 | Lung Carcinoma |
| MCF7 | PIK3CA, TP53, ESR1 | Breast Adenocarcinoma |
| HT29 | BRAF, TP53, APC | Colorectal Adenocarcinoma |
| HCT116 | KRAS, TP53, PIK3CA | Colorectal Carcinoma |
| HT1080 | NRAS, TP53 | Fibrosarcoma |
| HEPG2 | TP53, CTNNB1 | Hepatocellular Carcinoma (Liver) |
| U2OS | TP53, RB1 | Osteosarcoma (Bone) |

**Table S10.** Compounds selected by Prophet as potential MEK inhibitors, out of all 1.9M candidate compounds. Table containing the list of Prophet selected compounds. It identifies the compounds using ChEMBL ID and PubChem CID. It contains whether the compounds had available vendors and whether there exists prior literature relating the compound to MEK inhibition.

| **ChEMBL ID** | **PubChem CID** | **Vendor Available** | **MEK Literature** |
| --- | --- | --- | --- |
| chembl105442 | 6918454 | TRUE | TRUE |
| chembl1171312 | 25186684 | TRUE | TRUE |
| chembl1236682 | 44182295 | TRUE | TRUE |
| chembl1614701 | 10127622 | TRUE | TRUE |
| chembl1614766 | 11548630 | TRUE | TRUE |
| chembl1615025 | 24963252 | TRUE | TRUE |
| chembl2103875 | 11707110 | TRUE | TRUE |
| chembl2105684 | 16214875 | TRUE | TRUE |
| chembl2105741 | 50992434 | TRUE | TRUE |
| chembl2107832 | 44187362 | TRUE | TRUE |
| chembl2146883 | 16222096 | TRUE | TRUE |
| chembl2364607 | 71491931 | TRUE | TRUE |
| chembl244488 | 9803891 | TRUE | TRUE |
| chembl3182621 | 16666708 | TRUE | TRUE |
| chembl3187723 | 10288191 | TRUE | TRUE |
| chembl3330650 | 42642654 | TRUE | TRUE |
| chembl4297676 | 52918382 | TRUE | TRUE |
| chembl442235 | 11338183 | TRUE | TRUE |
| chembl447345 | 10117717 | TRUE | TRUE |
| chembl450775 | 10206158 | TRUE | TRUE |
| chembl481949 | 10112191 | TRUE | TRUE |
| chembl485964 | 5287529 | TRUE | TRUE |
| chembl507361 | 9826528 | TRUE | TRUE |
| chembl507836 | 10152076 | TRUE | TRUE |
| chembl5095241 | 71621329 | TRUE | TRUE |
| chembl510244 | 9953168 | TRUE | TRUE |
| chembl573579 | 45481530 | FALSE | FALSE |
| chembl573819 | 25163987 | TRUE | TRUE |
| chembl1169631 | 25187544 | FALSE | TRUE |
| chembl1169746 | 49797945 | FALSE | TRUE |
| chembl1169813 | 49797967 | FALSE | TRUE |
| chembl1169814 | 24784164 | FALSE | TRUE |
| chembl1169996 | 25186685 | FALSE | TRUE |
| chembl1169998 | 25187248 | FALSE | TRUE |
| chembl1169999 | 25186961 | FALSE | TRUE |
| chembl1170191 | 25187543 | FALSE | TRUE |
| chembl1170596 | 49797965 | FALSE | TRUE |
| chembl1170802 | 49797966 | FALSE | TRUE |
| chembl1170804 | 25186686 | FALSE | TRUE |
| chembl1170805 | 25186687 | FALSE | TRUE |
| chembl1171012 | 25186963 | FALSE | TRUE |
| chembl1171013 | 25186960 | FALSE | TRUE |
| chembl1171014 | 49798006 | FALSE | TRUE |
| chembl1171015 | 25188088 | FALSE | TRUE |
| chembl1171682 | 25186959 | FALSE | TRUE |
| chembl1172521 | 25187538 | FALSE | TRUE |
| chembl234887 | 11213014 | FALSE | TRUE |
| chembl234889 | 11269976 | FALSE | TRUE |
| chembl234890 | 11166713 | FALSE | TRUE |
| chembl235778 | 11213013 | FALSE | TRUE |
| chembl237256 | 11429515 | FALSE | TRUE |
| chembl237257 | 11165914 | FALSE | TRUE |
| chembl237258 | 11385135 | FALSE | TRUE |
| chembl237469 | 23630906 | FALSE | TRUE |
| chembl387574 | 44428116 | FALSE | TRUE |
| chembl391135 | 11464600 | FALSE | TRUE |
| chembl393282 | 11177427 | FALSE | TRUE |
| chembl395231 | 11189634 | FALSE | TRUE |
| chembl396548 | 44428115 | FALSE | TRUE |
| chembl396631 | 11406147 | FALSE | TRUE |
| chembl396632 | 11408896 | FALSE | TRUE |
| chembl397811 | 11362025 | FALSE | TRUE |
| chembl397812 | 11372988 | FALSE | TRUE |
| chembl442641 | 9954487 | FALSE | TRUE |
| chembl452600 | 10162188 | FALSE | TRUE |
| chembl453890 | 9804179 | FALSE | TRUE |
| chembl454114 | 9893385 | FALSE | TRUE |
| chembl454115 | 9872665 | FALSE | TRUE |
| chembl454116 | 9959922 | FALSE | TRUE |
| chembl454880 | 9824802 | FALSE | TRUE |
| chembl455470 | 22635332 | FALSE | TRUE |
| chembl455471 | 9803263 | FALSE | TRUE |
| chembl455472 | 9868895 | FALSE | TRUE |
| chembl464238 | 11533791 | FALSE | TRUE |
| chembl464239 | 11307224 | FALSE | TRUE |
| chembl464257 | 44570551 | FALSE | FALSE |
| chembl464258 | 44570552 | FALSE | FALSE |
| chembl465465 | 44570553 | FALSE | FALSE |
| chembl481968 | 44570599 | FALSE | FALSE |
| chembl481969 | 44570600 | FALSE | FALSE |
| chembl485736 | 10344482 | FALSE | TRUE |
| chembl485768 | 44589732 | FALSE | TRUE |
| chembl485944 | 10345414 | FALSE | TRUE |
| chembl485945 | 25023709 | FALSE | TRUE |
| chembl485951 | 11755808 | FALSE | TRUE |
| chembl485965 | 9983588 | FALSE | TRUE |
| chembl486138 | 10390808 | FALSE | TRUE |
| chembl486354 | 10458777 | FALSE | TRUE |
| chembl486559 | 11756341 | FALSE | TRUE |
| chembl487364 | 10455480 | FALSE | TRUE |
| chembl508526 | 22271652 | FALSE | TRUE |
| chembl510625 | 9959265 | FALSE | TRUE |
| chembl517647 | 44570554 | FALSE | FALSE |
| chembl518248 | 44570518 | FALSE | FALSE |
| chembl519555 | 10413260 | FALSE | TRUE |
| chembl521194 | 44589731 | FALSE | TRUE |
| chembl521370 | 11755809 | FALSE | TRUE |
| chembl521516 | 11754726 | FALSE | TRUE |
| chembl540501 | 45481529 | FALSE | FALSE |
| chembl4228999 | 137628642 | TRUE | TRUE |
| chembl3746640 | 51347355 | TRUE | TRUE |
| chembl3746559 | 127038617 | TRUE | TRUE |
| chembl2087082 | 44603148 | TRUE | TRUE |
| chembl519419 | 9887630 | TRUE | TRUE |
| chembl2146885 | 22025663 | TRUE | TRUE |
| chembl511056 | 9864956 | TRUE | TRUE |
| chembl3330652 | 44137226 | TRUE | TRUE |
| chembl3330649 | 25013308 | TRUE | TRUE |
| chembl2087078 | 46926364 | TRUE | TRUE |
| chembl2087460 | 44603207 | TRUE | TRUE |
| chembl2146887 | 16223325 | TRUE | TRUE |
| chembl2146888 | 16222191 | TRUE | TRUE |
| chembl2146886 | 16223412 | TRUE | TRUE |
| chembl3645238 | 70997213 | TRUE | FALSE |
| chembl3590107 | 24866313 | TRUE | TRUE |
| chembl3335206 | 17102070 | TRUE | FALSE |
| chembl1474985 | 9956637 | TRUE | TRUE |
| chembl4127114 | 145439085 | TRUE | TRUE |
| chembl3819302 | 127052212 | TRUE | TRUE |
| chembl2131626 | 67065878 | TRUE | FALSE |
| chembl1684070 | 24963251 | TRUE | TRUE |
| chembl1684063 | 53323815 | TRUE | TRUE |
| chembl3349342 | 15985330 | TRUE | TRUE |

**Table S11.** Vendors from where the compounds were bought.

| ChEMBL identifier | Common Name | Vendor | Vendor identifier |
| --- | --- | --- | --- |
| chembl450775 | PD-0325901,<br>Mirdametinib | KeyOrganics | PD0325901 |
| chembl244488 | PD-0316684 | KeyOrganics | AS-73133 |
| chembl3335206 |  | ChemBridge | SC-9110770 |
| chembl3645238 | SCH772984 | MCULE | MCULE-2500426897 |
| chembl507836 |  | ChemDIV | Y700-0482 |
|  | Staurosporine | Santa Cruz<br>Biotechnology | sc-3510A |

